# Acute myeloid leukemia mitochondria hydrolyze ATP to resist chemotherapy

**DOI:** 10.1101/2024.04.12.589110

**Authors:** James T Hagen, Mclane M Montgomery, Raphael T Aruleba, Brett R Chrest, Thomas D Green, Miki Kassai, Tonya N Zeczycki, Cameron A Schmidt, Debajit Bhowmick, Su-Fern Tan, David J Feith, Charles E Chalfant, Thomas P Loughran, Darla Liles, Mark D Minden, Aaron D Schimmer, Myles C Cabot, Joseph M Mclung, Kelsey H Fisher-Wellman

## Abstract

Despite early optimism, therapeutics targeting oxidative phosphorylation (OxPhos) have faced clinical setbacks, stemming from their inability to distinguish healthy from cancerous mitochondria. Herein, we describe an actionable bioenergetic mechanism unique to cancerous mitochondria inside acute myeloid leukemia (AML) cells. Unlike healthy cells which couple respiration to the synthesis of ATP, AML mitochondria were discovered to support inner membrane polarization by consuming ATP. Because matrix ATP consumption allows cells to survive bioenergetic stress, we hypothesized that AML cells may resist cell death induced by OxPhos damaging chemotherapy by reversing the ATP synthase reaction. In support of this, targeted inhibition of BCL-2 with venetoclax abolished OxPhos flux without impacting mitochondrial membrane potential. In surviving AML cells, sustained polarization of the mitochondrial inner membrane was dependent on matrix ATP consumption. Mitochondrial ATP consumption was further enhanced in AML cells made refractory to venetoclax, consequential to downregulations in both the proton-pumping respiratory complexes, as well as the endogenous F_1_-ATPase inhibitor *ATP5IF1*. In treatment-naive AML, *ATP5IF1* knockdown was sufficient to drive venetoclax resistance, while *ATP5IF1* overexpression impaired F_1_-ATPase activity and heightened sensitivity to venetoclax. Collectively, our data identify matrix ATP consumption as a cancer-cell intrinsic bioenergetic vulnerability actionable in the context of mitochondrial damaging chemotherapy.

## INTRODUCTION

Similar to a growing list of human solid organ malignancies, AML is particularly reliant on oxidative metabolism^1–7^. Pioneering studies over the past several years have repeatedly demonstrated potent anti-leukemic effects upon targeting AML’s mitochondrial reliance^8–15^. Although mitochondrial targeted therapies are entering clinical trials, the majority of hopeful ‘mito-therapeutics’ suffer from a limited therapeutic index based on their inability to discriminate cancerous from non-cancerous mitochondria^11,16–18^. The lack of cancer-specific mitochondrial targets poses a significant barrier to progress, as safe and effective mitochondrial-targeted therapies will undoubtedly require cancer cell selectivity. Fortunately, new evidence is emerging that all mitochondria are not alike. In fact, mitochondria are unique in composition and function within each cell type, including AML cells^19–22^. This raises the exciting possibility that identifying the unique bioenergetic signature(s) of AML cells may hold the key to designing mitochondrial-targeted therapies that preferentially eradicate the malignant cells.

Compared to normal blood cells, AML cells present with intrinsic deficiencies in OxPhos^22^. Despite AML cells having a larger mitochondrial network, the fraction of this network capable of coupling respiration to the synthesis of ATP is less than 50%^22^. Importantly, this fraction has been found to fall below 30% in chemoresistant AML^23^. Thus, although oxidative ATP synthesis is essential for most all eukaryotic life, there is emerging evidence linking impaired OxPhos function to AML disease progression^22,23^. With respect to chemoresistant disease, despite intrinsic OxPhos limitations, mitochondrial membrane potential ( ΔΨ_m_) is either sustain or hyperpolarized in drug resistant AML cells^3,24^. Because mitochondrial hyperpolarization typically tracks with high OxPhos, the low OxPhos yet hyperpolarized phenotype of drug resistant AML cells is unexpected and as such, highly diagnostic of cell-intrinsic mitochondrial remodeling^25,26^.

Mitochondrial polarization is canonically generated via the proton pumping respiratory complexes (i.e., complexes I/III/IV). In respiratory incompetent cells, polarization can also be sustained via cytosol to matrix ATP transport and/or ATP hydrolysis by the reversible ATP synthase reaction^27–31^. Thus, in the absence of a sufficient proton-motive force, ATP synthase reverts to an F_1_-ATPase, converting the free energy available in the cell’s phosphate potential to a polarized mitochondrial inner membrane^27–31^. Maintenance of ΔΨ_m_ via matrix F_1_-ATPase activity has been demonstrated to be critical for cell survival in the context of respiratory complex lesions^27,32^. Mitochondrial depolarization is a key trigger for apoptosis^33,34^, therefore mitochondrial ATP consumption may also limit the cytotoxicity of AML cells in response to OxPhos damaging chemotherapy.

Designed to directly initiate the intrinsic pathway of apoptosis at the mitochondria^35^, the targeted therapy venetoclax (i.e., a Bcl-2 inhibitor and a mediator of BAX/BAK dependent apoptosis) was granted regular FDA approval for the treatment of AML in 2020^36^. Despite improving patient outcomes, clinical efficacy of venetoclax remains stifled by the development of chemoresistant disease^37,38,39^. Prior research has established that venetoclax exposure inhibits AML mitochondrial respiration^40^ by impinging on components of the electron transport system (ETS)^41^. Although respiration is inhibited, AML cells exposed to venetoclax sustain polarization across the mitochondrial inner membrane^42,43^. How AML cells sustain ΔΨ_m_ in the presence of venetoclax, and whether sustained ΔΨ_m_ drives venetoclax resistance, remains unclear. To investigate this, we leveraged an in-house mitochondrial phenotyping platform^44^ to systematically define the interplay between venetoclax exposure, mitochondrial bioenergetics, and AML survival.

## RESULTS

### Intrinsic lesions in respiratory complex IV limit OxPhos flux in AML cells

Relative to normal myeloid progenitor cells, AML blasts are characterized by high mitochondrial content^6,22^. Despite having more mitochondria per cell, we previously demonstrated that less than half of the AML mitochondrial network is capable of coupling respiration to the synthesis of ATP – a phenomenon indicative of OxPhos limitations^22,23^. Across a panel of AML cells lines (both human^22,23^ and mouse^19^), as well as AML patient samples, 50% or less of total respiratory flux (“Respiratory Capacity”) was stimulated during OxPhos (**Fig. 1A**) (**Supp. Fig 1A-H**). With that percentage below 40%, OxPhos limitations are even more pronounced in cells made resistant to chemotherapy^23^ (**Fig. 1A**), suggesting that intrinsic limitations in OxPhos flux is a conserved bioenergetic feature of AML cells with potential implications for combating treatment resistance.

**Figure 1.**
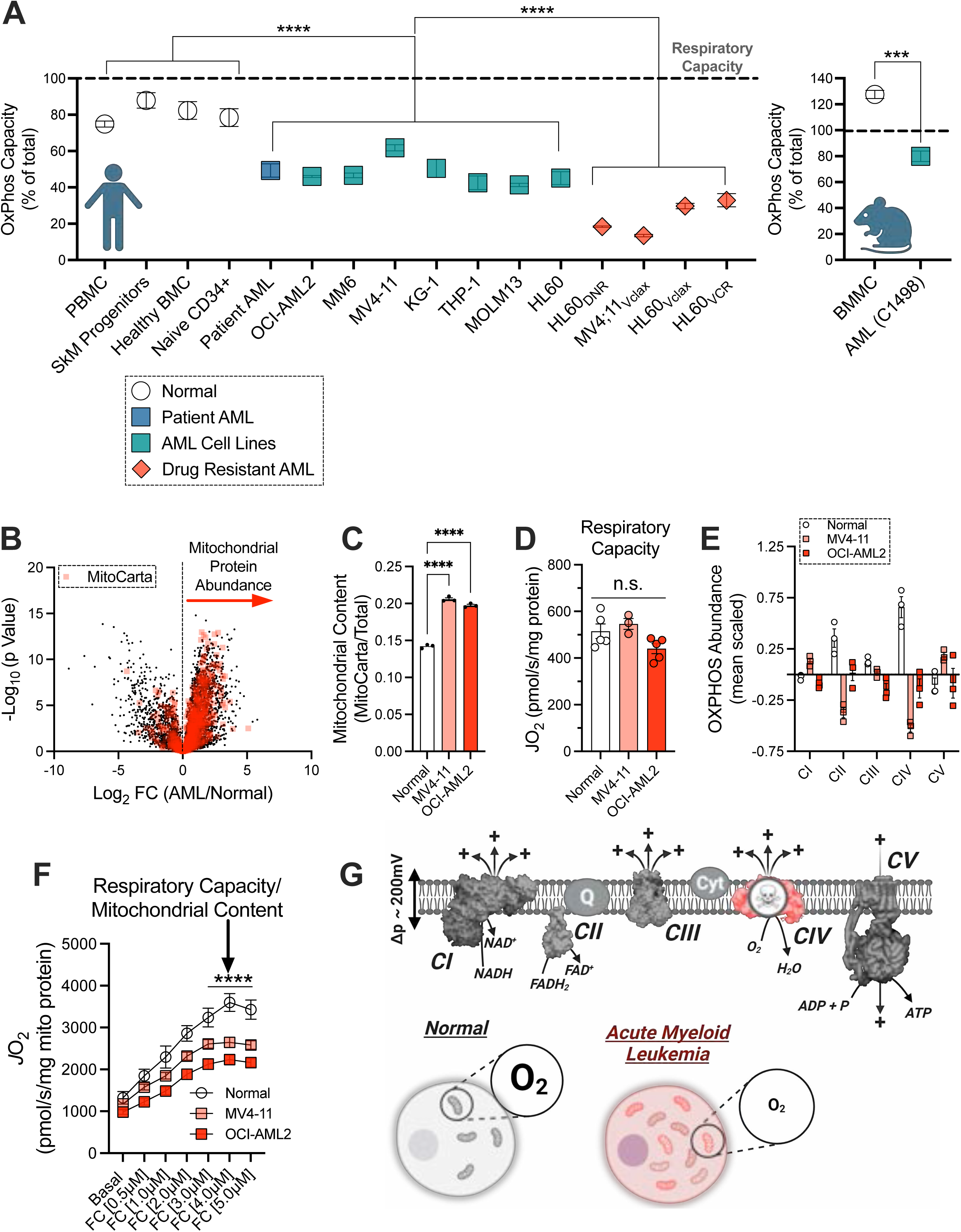
Intrinsic lesions in respiratory complex IV limit OxPhos flux in AML cells. All experiments were performed using whole cells and digitonin-permeabilized cells (A) Comparison of fractional OXPHOS of normal blood cells, patient AML, AML cell lines, chemoresistant AML, and mouse AML calculated as the ratio of *J*H^+^_OXPHOS_ to *J*H^+^_Total_ (*n* = 3-30 replicates) and represented as a percentage of total respiratory capacity. SkM Progenitors refers to myoblasts isolated from human skeletal muscle biopsy. (B) Volcano plot comparing mitochondrial and non-mitochondrial proteome of AML cells and healthy bone marrow mononuclear cells (Healthy BMC) (C) Comparison of mitochondrial content in AML cells and Healthy BMC cells (*n* = 3-30 replicates). (D) Comparison of whole cell respiratory capacity in Healthy BMC, MV4-11 and OCI-AML2 cells normalized to milligrams of protein (*n* = 3-5 replicates). (E) Comparison of the OXPHOS proteome between Healthy BMC and AML cells (*n* = 3 replicates). (F) Comparison of whole cell respiratory capacity in Healthy BMC, MV4-11 and OCI-AML2 cells normalized to milligrams of mitochondrial protein (*n* = 3-5 replicates). (G) Schematic depicting complex IV lesions and reduced respiration of individual AML mitochondria. Data are presented as mean ±SEM and analyzed by two-way ANOVA (E,F) and one-way ANOVA (A,C,D). *p<0.05, **p<0.01, ***p<0.001, ****p<0.0001.

To explore the underlying mechanisms of OxPhos limitations in AML we compared OxPhos complex expression and OxPhos respiratory flux between AML cell lines (e.g., MV4-11 and OCI-AML2) and healthy myeloid progenitors isolated from bone marrow aspirates. As reported previously^6,22^, mitochondrial protein abundance was higher in AML blasts, corresponding to a ∼ 40% increase in mitochondrial content (**Fig. 1B-C**). High mitochondrial content would be expected to lead to a corresponding increase in cellular respiratory capacity. However, protein normalized respiratory capacity was not elevated in AML blasts (**Fig. 1D**), indicating discordance between mitochondrial network size and respiratory kinetics – a feature diagnostic of respiratory complex lesions. To investigate the intrinsic composition of the respiratory system in AML, the expression of each respiratory complex and ATP synthase (i.e., the OxPhos components) were normalized to mitochondrial content. In doing so, mean scaled abundance of several respiratory complexes were observed to be lower in AML cells (**Fig. 1E**) (**Supp. Fig. 1I-M**). Complex specific lesions were most pronounced for cytochrome oxidase (i.e., complex IV), the terminal O_2_ consuming enzyme in the ETS (**Fig. 1E**). Consistent with reduced ETS expression, cellular respiratory capacity per unit of mitochondrial protein was lower in AML cells (**Fig. 1F**). These data agree with recent findings in mouse AML cells where we had observed AML-specific lesions in complexes III and IV sufficient to reduce OxPhos flux^19^. Together, these data implicate reduced OxPhos function as a conserved bioenergetic feature of AML cells driven by intrinsic lesions in the proton pumping respiratory complexes. AML cells appear circumvent these ETS lesions to sustain baseline respiratory flux by expanding mitochondrial content (**Fig. 1G**).

### Mitochondria inside AML cells consume cellular ATP to contribute to mitochondrial polarization

Despite deficiencies in OxPhos, AML cells have more mitochondria than normal cells and those mitochondria generate a voltage potential^22,45^. The ΔΨ_m_ can become polarized in two different ways: 1) via the proton-pumping respiratory complexes – complexes I/III/IV; or 2) matrix ATP uptake via the adenine nucleotide translocase (ANT) and/or hydrolysis of ATP by the F_1_-ATPase^46^. Because AML cells present with intrinsic respiratory complex lesions that would be expected to weaken the voltage generated by the ETS, we hypothesized that polarization may be supported via F_1_-ATPase activity. To test this, we used flow cytometry to assess mitochondrial polarization (using TMRM fluorescence) in individual AML cells in the absence and presence of the ATP synthase inhibitor oligomycin. Operating in non-quench mode, an increase in TMRM fluorescence with oligomycin was taken to indicate functional OxPhos, whereas a decrease in TMRM fluorescence with oligomycin was assumed to indicate ΔΨ_m_ was being generated, in part, via F_1_-ATPase activity (**Fig. 2A**). To validate our assay, ΔΨ_m_ was assessed in respiration-incompetent p^0^ OCI-AML2 (“OCI_Rho0_”) cells^47^ that present with a lack of mitochondrial respiration and a hyper-reduced NADH pool. (**Supp. Fig. 2A-B**). Despite being depleted of mtDNA, p^0^ cells sustain ΔΨ_m_ by consuming ATP^27^. In agreement, inhibition of ATP synthase with oligomycin triggered depolarization in OCI_p0_ cells (**Fig. 2B-C**). In mononuclear cells isolated from healthy bone marrow, as well as PBMCs from healthy volunteers, the addition of oligomycin increased TMRM fluorescence, indicating functional OxPhos (**Fig. 2D-E**). Interestingly, ΔΨ_m_ was sustained in the presence of rotenone (i.e., a Complex I inhibitor) and antimycin A (i.e., an inhibitor of Complex III), indicating that normal blood cells can sustain ΔΨ_m_ via matrix ATP consumption when respiration is compromised (**Fig. 2D-E**).

**Figure 2.**
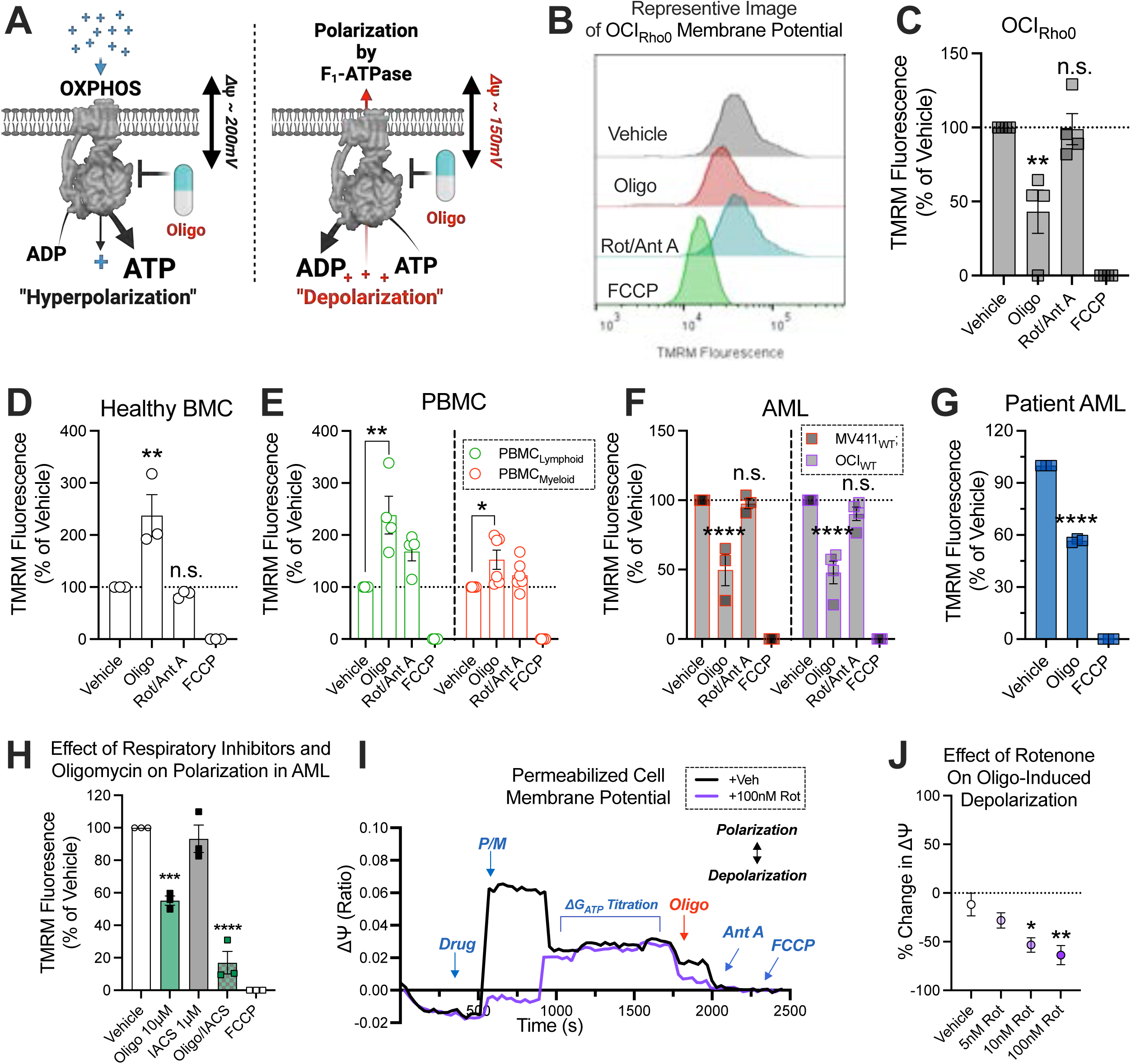
Mitochondria in AML cells consume cellular ATP to sustain mitochondrial polarization. All experiments were performed using whole intact cells. (A) Schematic depicting strategy to determine functional OxPhos in ΔΨ_m_ assays. (B) Representative image from flow cytometric analysis of intact cell ΔΨ_m_ in OCI_Rho0_ cells. (C) Flow cytometric analysis of intact cell ΔΨ_m_ in OCI_Rho0_ cells (*n* = 4 replicates). (D) Flow cytometric analysis of intact cell ΔΨ_m_ in Healthy BMC (*n* = 3 replicates). (E) Flow cytometric analysis of intact cell ΔΨ_m_ in myeloid and lymphoid populations sorted from PBMC (*n* = 4-6 replicates). (F) Flow cytometric analysis of intact cell ΔΨ_m_ in AML cell lines (*n* = 3-4 replicates). (G). Flow cytometric analysis of intact cell ΔΨ_m_ in patient AML (*n* = 3 replicates). (H) Flow cytometric analysis of intact cell ΔΨ_m_ in pooled MV411_WT_, HL60_WT_, and OCI_WT_ cells (*n* = 1 replicate per cell type). (I) Representative trace of permeabilized cell ΔΨ_m_ assay in OCI_WT_ cells. (J) Comparison of oligomycin-induced depolarization in OCI_WT_ cells in the presence of increasing doses of rotenone (*n* = 4 replicates). Data are presented as mean ±SEM and analyzed by two-way ANOVA (F), one-way ANOVA (C,D,G,H,J), or unpaired t-test (E). *p<0.05, **p<0.01, ***p<0.001, ****p<0.0001.

Having established the validity of our approach, we then assessed ΔΨ_m_ in AML cells. Across two AML cell lines with different founding mutations (MV4-11, “MV411_WT_” and OCI-AML2, “OCI_WT_”), as well as patient AML samples, the addition of oligomycin lowered TMRM fluorescence (**Fig. 2F-G**). Remarkably, the magnitude of depolarization in AML cells upon ATP synthase inhibition was comparable to that seen in respiration deficient OCI_p0_ cells. (**Fig. 2C) (Fig 2F-G**). Using both oligomycin to block ATP synthase and IACS-010759 to block the ETS near-complete depolarization was achieved (**Fig 2H**), suggesting both the ETS and ATP consumption cooperate to sustain ΔΨ_m_. To control for effects driven by constituents in IMDM culture media, ΔΨ_m_ in OCI_WT_ cells was assessed in a culture medium formulated to mimic the physiological profile of human plasma (i.e. Human Plasma-Like Medium, or HPLM). Oligomycin also triggered depolarization in OCI_WT_ cells assayed in HPLM (**Supp. Fig. 2C**). To validate these data with an alternative approach, fluorescent microscopy was used to measure changes in TMRM fluorescence (operating in non-quench mode) of MV411_WT_ cells in response to either oligomycin, rotenone and antimycin A, or those in combination. Single agent oligomycin depolarized ΔΨ_m_ in MV411_WT_ cells (**Supp. Fig. 2D-E**). Rotenone and antimycin A also resulted in depolarization, however, when oligomycin, rotenone and antimycin A were used in combination, the magnitude of depolarization was similar to that of alamethicin -- a channel forming peptide antibiotic that eliminates ΔΨ_m_ (**Supp. Fig. 2D-E**).

The consumption of ATP by AML mitochondria appears consequential of a weakened voltage potential driven in part, by intrinsic lesions in the respiratory complexes. We hypothesized that mimicking complex specific lesions would increase the reliance on F_1_-ATPase activity to sustain polarization. To test this, we designed an assay that measures ΔΨ_m_ of digitonin-permeabilized AML cells by quantifying changes in TMRM fluorescence operating in quench mode. To begin, OCI_WT_ mitochondria were energized with Complex I substrates (i.e., pyruvate and malate, “P/M”) and exposed to increasing doses of the complex I inhibitor rotenone. The ΔΨ_m_ was measured across a span of ATP/ADP ratios with ΔG_ATP_ values ranging from -54.2 kJ/mol to -61.5 kJ/mol. Following ΔG_ATP_ titration, oligomycin was added to inhibit F_1_-ATPase activity and changes in TMRM fluorescence were quantified. A dose-dependent inhibition of complex I-driven polarization was induced by rotenone and that tracked with a larger magnitude of depolarization by oligomycin, suggesting that complex-specific lesions enhance the reliance on F_1_-ATPase activity to sustain polarization in AML cells (**Fig. 2I-J**) (**Supp. Fig. 2F**). Regarding the source of ATP being consumed by AML cells, we considered contributions by both glycolysis^48^ (i.e., in the cytosol) or glutaminolysis^49^ (i.e., in the matrix). To test, the depolarizing effects of oligomycin were quantified in OCI_WT_ cells in the presence of either 2-DG (i.e., an inhibitor of glycolysis) or CB-839 (i.e., an inhibitor of glutaminase). Despite inhibiting either glycolysis or glutaminolysis, the magnitude of depolarization induced by oligomycin was not impacted (**Supp. Fig. 2G**). These data suggest that ATP fueling F_1_-ATPase activity may not be confined to only one source. Collectively, our data demonstrate that unlike normal blood cells with functional OxPhos, AML mitochondria consume ATP to support polarization of the ΔΨ_m_.

### Matrix ATP consumption via F_1_-ATPase activity is an actionable vulnerability in AML cells

Mitochondria in AML cells appear to respire and simultaneously reverse the ATP synthase reaction. However, when F_1_- ATPase activity was inhibited by oligomycin, the ETS was unable to re-polarize the inner membrane back to baseline. These data are surprising, as they suggest that upon inhibition of the F_1_-ATPase in AML, polarization originating from the ETS must also become inhibited. Inhibition of respiration by matrix ATP is a phenotype of AML cells that we have reported previously^22^. Expanding on those findings, when the ETS was directly accessed by permeabilizing MV411_WT_ mitochondria using alamethicin, respiration supported by Complex I (NADH) and Complex II (succinate) substrates was dose-dependently inhibited by ΔG_ATP_ (**Fig. 3A**) (**Supp. Fig. 3A-B**). These data indicate that AML respiration is directly inhibited by matrix ATP. Because matrix ATP is consumed by the F_1_-ATPase, we hypothesized that F_1_-ATPase activity supports AML respiration by clearing matrix ATP. To test this, digitonin-permeabilized OCI_WT_ cells were energized with saturating carbon substrates (pyruvate, malate, glutamate, succinate, and octanoyl-carnitine; “P/M/G/S/O”) and respiration was stimulated using FCCP. Cells were then provided DMSO or oligomycin and ΔG_ATP_ was titrated to inhibit respiration. In the presence of oligomycin, OCI_WT_ cell respiration was more sensitive to inhibition by ΔG_ATP_, indicating that F_1_- ATPase activity can support respiration by degrading matrix ATP (**Fig. 3B**). To determine if respiration in living AML cells also becomes inhibited when F_1_-ATPase is blocked, intact OCI_WT_ cell respiration was stimulated using FCCP and then either oligomycin or no oligomycin was added to inhibit F_1_-ATPase activity. Oligomycin inhibited uncoupled respiration in intact OCI_WT_ cells, suggesting that F_1_-ATPase activity supports respiration in living AML cells (**Fig. 3C**).

**Figure 3.**
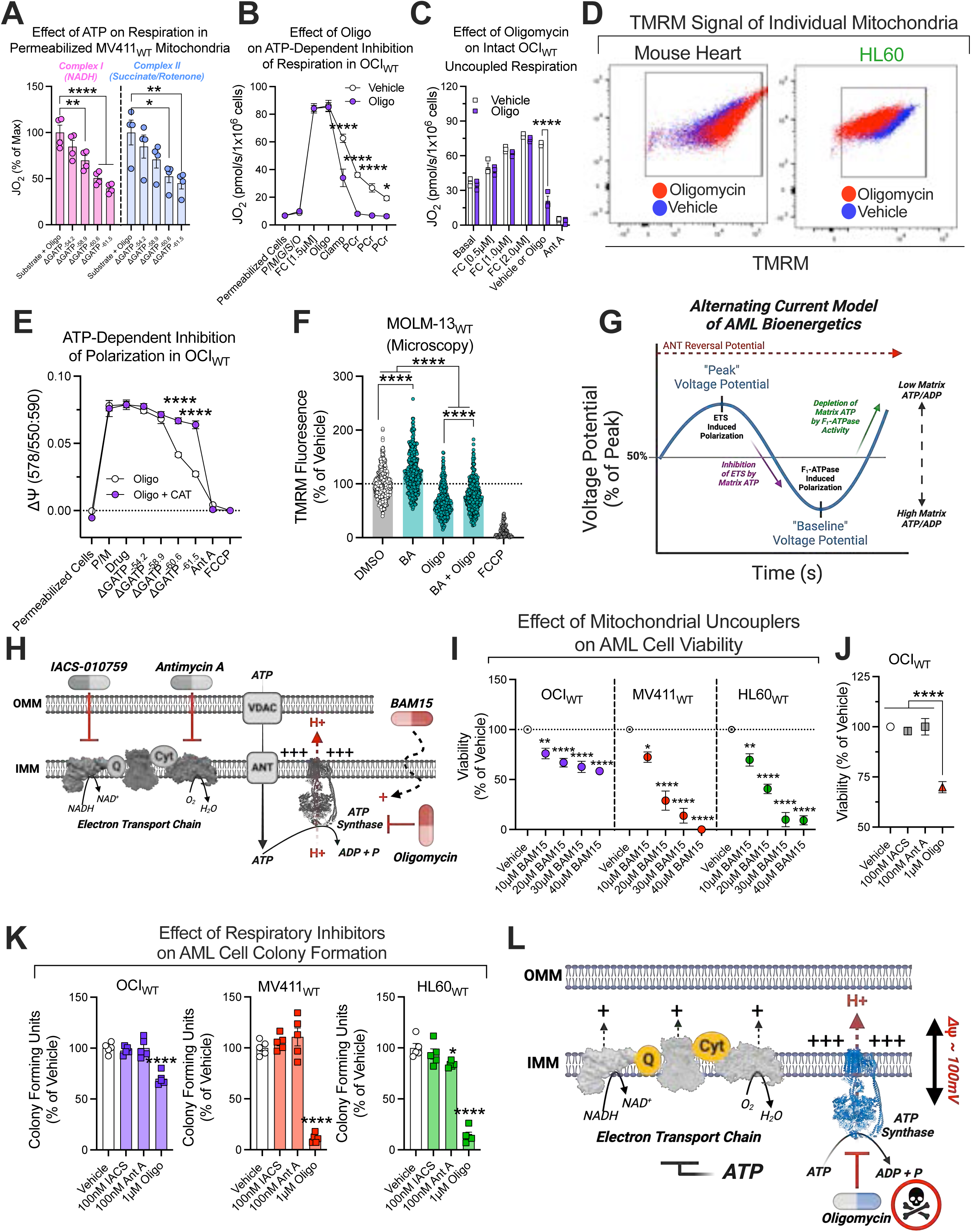
F_1_-ATPase activity is an actionable vulnerability of AML cells. All experiments were performed using whole intact cells, digitonin-permeabilized cells, or alamethicin-permeabilized mitochondria. (A) Effect of ΔG_ATP_ on maximal NADH (Complex I) or succinate/rotenone (Complex II) supported respiration in alamethicin-permeabilized MV411_WT_ mitochondria (*n* = 4 replicates). (B) Effect of oligomycin on inhibition of uncoupled respiration by ΔG_ATP_ in permeabilized OCI_WT_ cells (*n* = 3 replicates). (C) Effect of oligomycin on uncoupled respiration in intact OCI_WT_ cells (*n* = 3 replicates). (D) Change in TMRM fluorescence of individual mitochondria in whole mitochondrial populations isolated from mouse heart or HL60 before and after oligomycin addition during flow cytometric analysis of the ΔΨ_m_. (E) Effect of ΔG_ATP_ on polarization by P/M/G/S/O in the presence of oligomycin, and in the presence or absence of carboxyatractyloside, in permeabilized OCI_WT_ cells (*n* = 3 replicates). (F) Fluorescent microscopy analysis of intact cell ΔΨ_m_ in MOLM-13_WT_ cells in the presence bongkrekic acid, oligomycin, or those in combination (*n* = 189-417 cells). (G) Schematic depicting an alternating current model of AML mitochondrial bioenergetics. (H) Schematic depicting the mitochondrial targets of IACS, antimycin A, oligomycin, and BAM15 uncoupler. (I) Effect of increasing doses of BAM15 uncoupler on OCI_WT_, MV411_WT_, and HL60_WT_ cell viability. Viability was measured using propidium iodide (*n* = 4-5 replicates). (J) Effect of IACS, antimycin A, or oligomycin on OCI_WT_ cell viability. Viability was measured using propidium iodide (*n* = 5 replicates). (K) Effect of IACS, antimycin A, or oligomycin on OCI_WT_, MV411_WT_, and HL60_WT_ colony formation. Colonies were manually counted using microscopy (*n* = 5 replicates). (L) Schematic depicting mechanism of AML cell death by oligomycin. Data are presented as mean ±SEM and analyzed by two-way ANOVA (A,C,E,F) or one-way ANOVA (A,I,J,K). *p<0.05, **p<0.01, ***p<0.001, ****p<0.0001.

It is possible that individual AML cells contain two seperate mitochondrial populations - a population that respires and synthesizes ATP, and a population that cannot respire but consumes ATP. To test this, we designed an assay that measures ΔΨ_m_ (using TMRM fluorescence operating in non-quench mode) of individual mitochondria. To begin, mitochondria isolated from mouse heart or HL60 cells were energized with pyruvate/malate and a ΔG_ATP_ value of -61.5 kJ/mol, followed by the addition of oligomycin. Oligomycin increased polarization in mouse heart, and decreased polarization in HL60 cells (**Supp. Fig. 3C-D**). TMRM fluorescence of individual mitochondria within the mouse heart or HL60 mitochondrial population were plotted before (“Vehicle”) and after the addition of oligomycin (“Oligomycin”) and appeared to shift homogenously (**Fig. 3D**) (**Supp. Fig. 3D**). These data suggest that AML cells contain a single mitochondrial population that simultaneously respires and consumes ATP.

Given that matrix ATP inhibits AML mitochondrial respiration, we hypothesized that matrix ATP may also blunt ETS polarization. To test this, we measured the effect of ATP on ΔΨ_m_ of permeabilized OCI_WT_ cells polarized with pyruvate/malate in the presence of oligomycin to inhibit the F_1_-ATPase. OCI_WT_ cells were provided either no carboxyatractyloside (“CAT”) or CAT to inhibit ANT-mediated matrix ATP uptake, and ΔG_ATP_ was then titrated. The ΔG_ATP_ titration triggered depolarization of OCI_WT_ cells which was rescued by CAT (**Fig. 3E**). These data indicate that respiratory inhibition by matrix ATP inhibits ETS polarization and suggests that ΔΨ_m_ in AML cells is insufficient to fuel ANT-mediated matrix to cytosol ATP transport ^31^. Thus, matrix ATP clearance by the F_1_-ATPase may be necessary to prevent matrix ATP from blunting respiration and polarization by the ETS in AML cells. In living AML cells, we hypothesized that inhibition of ANT with bongkrekic acid (i.e., “BA”, a cell-permeable inhibitor of ANT) would rescue oligomycin-induced depolarization by blocking cytosol to matrix ATP transport. To test this, we quantified the ΔΨ_m_ in living MOLM-13_WT_ cells in the presence of either BA, oligomycin, or that in combination. Although BA increased polarization in the presence and absence of oligomycin, BA was insufficient to rescue depolarization by oligomycin (**Fig. 3F**) (**Supp. Fig. 3E**). However, when we measured the effect of BA on ΔΨ_m_ in living, respiration deficient OCI_p0_ cells, BA triggered depolarization (**Supp. Fig. 3F**). These data suggest that respiration deficient OCI_Rp0_ mitochondria consume cytosolically derived ATP to sustain polarization. In AML cells that do respire, some matrix ATP may be derived from the cytosol, however, a portion of ATP in the matrix space appears to be synthesized and degraded in the matrix by a reversible F_1_-ATPase/ATP synthase reaction, akin to a futile energy cycle (**Fig. 3G**). We suspect that the direction of the F_1_-ATPase/ATP synthase reaction in respiring AML cells is driven by fluctuations in polarization, thereby mimicking an alternating current circuit (**Fig. 3G**). Nonetheless, F_1_-ATPase activity in AML cells appears to be a mechanism necessary to degrade matrix ATP, regardless of the source of that ATP, thereby preventing depolarization.

Depolarization is a known driver of cell death via apoptosis^33,34^. When AML cells were exposed to increasing doses of BAM15, a depolarizing respiratory uncoupler, dose-dependent toxicity was achieved (**Fig. 3H-I**). Given that F_1_-ATPase activity is important for AML cells to remain polarized, we hypothesized that F_1_- ATPase activity is important for AML survival. To test this, AML cell viability and colony formation were quantified in response to either oligomycin, IACS, or antimycin A as single agents. Although IACS and antimycin A inhibited basal respiration (**Supp. Fig. 3G**), AML cells were resistant to overt losses in cell viability and colony formation in response to either drug (**Fig. 3H, J-K**), presumably because the cells could sustain polarization via ATP consumption. Consistent with this, oligomycin resulted in greater losses of colony formation and cell viability, indicating F_1_-ATPase activity is important for AML cell survival (**Fig. 3J-K**). Although AML cells can polarize via both the ETS and the F_1_-ATPase, our data suggest that oligomycin results in an inhibition of polarization originating from both the F_1_-ATPase (i.e., by inhibition of the F_1_-ATPase with oligomycin) and the ETS (i.e., by inhibition from matrix ATP) (**Fig. 3L**). Collectively, these data demonstrate that F_1_-ATPase activity is an actionable vulnerability of AML cells (**Fig. 3L**).

### Venetoclax exposure collapses mitochondrial respiration, but polarization is sustained via F_1_-ATPase activity

Inhibition of respiration by matrix ATP appears to be a conserved bioenergetic mechanism that, by depolarizing the inner membrane, facilitates the apoptosis of cells – particularly those with mitochondrial damage. For example, despite intrinsic lesions in complex IV, AML cells survive by degrading matrix ATP via F_1_-ATPase activity (**Fig 3J)**. By default, polarization via F_1_-ATPase activity also confers AML cells an inherent resistance to respiratory inhibition (**Fig. 3J-K**). Disruptions in AML cell respiration have been reported to occur in response to various chemotherapeutics including doxorubicin^50^, cytarabine^51^, and venetoclax^40^. Venetoclax, a specific inhibitor of the anti-apoptotic Bcl-2 protein, has been shown to inhibit respiratory flux in AML cells by impinging on the proton-pumping respiratory complexes^40,41^. In our hands, 1-hour exposure to venetoclax in treatment naïve AML cells (“MV411_WT_” and “OCI_WT_”) collapsed mitochondrial respiration (i.e., both respiratory capacity and OxPhos respiratory flux), without overtly reducing cell viability (**Fig. 4A-D**) (**Supp. Fig. 4A-B**). Consistent with venetoclax mediating BAX/BAK-dependent outer mitochondrial membrane permeabilization^35^, the respiration of AML cells exposed to venetoclax was increased following the addition of cytochrome C, indicating that Bcl-2 inhibition results in damage to the outer mitochondrial membrane (**Fig. 4C**). Mitochondria in living AML cells have been shown to remain polarized in response to venetoclax^24^. Because venetoclax collapsed mitochondrial respiration in AML cells, we hypothesized that polarization may be sustained via F_1_- ATPase activity. To test this, we leveraged our permeabilized cell ΔΨ_m_ assay. In this assay, polarization induced by various substrates was measured following 1-hour exposure of treatment naïve AML cells to venetoclax. Venetoclax exposure resulted in an inhibition of polarization by Complex I (pyruvate/malate/glutamate or “P/M/G”) and Complex II (succinate/rotenone or “S/Rot”) substrates (**Fig. 4E**) (**Supp. Fig. 4C-D**). However, polarization by F_1_-ATPase activity (“ATP”) was not overtly affected by venetoclax exposure (**Fig. 4E**) (**Supp. Fig. 4C-D**), suggesting F_1_-ATPase activity sustains polarization across the inner membrane during Bcl-2 inhibition.

**Figure 4.**
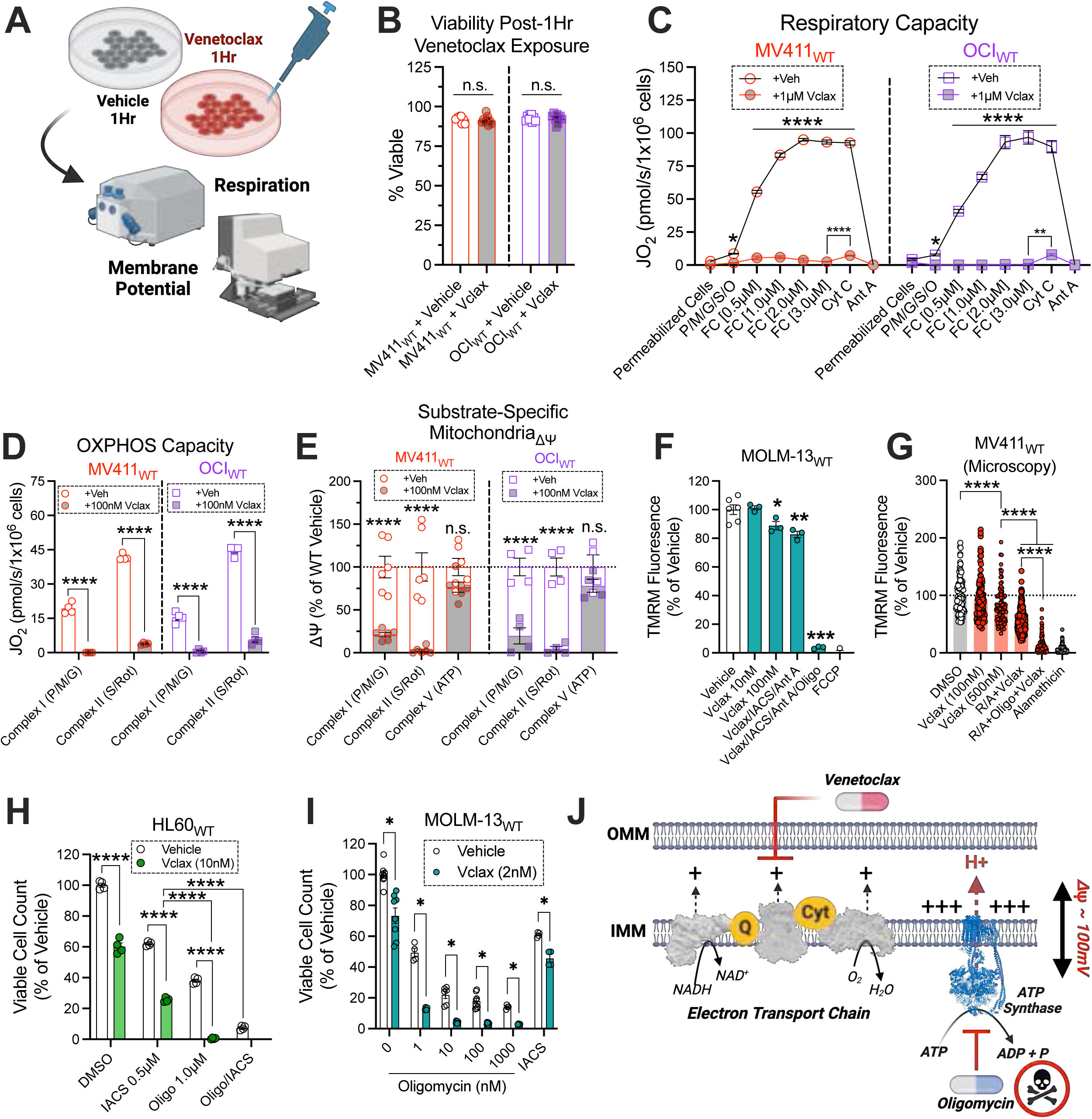
Venetoclax exposure collapses mitochondrial respiration, but polarization is sustained via F_1_- ATPase activity. All experiments were performed using whole intact cells or digitonin permeabilized cells. (A) Schematic depicting characterization of treatment naïve AML cells exposed to venetoclax for 1 hour. (B) Effect of 1 hour exposure to venetoclax on AML cell viability. Viability was measured using trypan blue viable cell count (*n* = 9 replicates). (C) Effect of 1 hour exposure to 1µM venetoclax on respiratory capacity in permeabilized AML cells (*n* = 3 replicates). (D) Effect of 1 hour exposure to 100nM venetoclax on OxPhos capacity in permeabilized AML cells (*n* = 4 replicates). (E) Effect of 1 hour exposure to 100nM venetoclax on polarization induced by Complex I substrates (P/M/G), Complex II substrates (S/R), or Complex V substrates (ATP) in permeabilized AML cells (*n* = 4-6 replicates). (F) Flow cytometric analysis of intact MOLM-13_WT_ cell ΔΨ_m_ in response to single agent venetoclax, and venetoclax in combination with IACS, antimycin A, and those in combination with oligomycin (*n* = 3 replicates). (G) Fluorescent microscopy analysis of intact MV411_WT_ cell ΔΨ_m_ in response to single agent venetoclax, and venetoclax in combination with rotenone, antimycin A, and those in combination with oligomycin (*n* = 100-311 cells). (H) Effect of single agent venetoclax, venetoclax in combination with IACS and oligomycin, or IACS in combination with oligomycin on HL60_WT_ cell viability. Viability was measured using CellTiter-Glo (*n* = 4 replicates). (I) Effect of single agent venetoclax or venetoclax in combination with either oligomycin or IACS on MOLM-13_WT_ cell viability. Viability was measured using CellTiter-Glo (*n* = 4-8 replicates). (J) Schematic depicting mechanism of AML cell death in response to venetoclax and oligomycin. Data are presented as mean ±SEM and analyzed by two-way ANOVA (C,D,E,G,H), one-way ANOVA (B,F), or unpaired t-test (I). Cytochrome C effect in (C) analyzed by unpaired t-test. *p<0.05, **p<0.01, ***p<0.001, ****p<0.0001.

Although venetoclax is an emerging frontline chemotherapeutic in AML, the development of chemoresistant disease limits the drug’s clinical efficacy^37,38,39^. In pre-clinical models, the efficacy of venetoclax has been improved by inhibiting respiration (e.g., with IACS) in combination with venetoclax^52,53^. During venetoclax exposure however, our findings indicate that AML cells sustain polarization via F_1_-ATPase activity (**Fig. 4E**) (**Supp. Fig. 4C-D**). Similarly, F_1_-ATPase activity sustains polarization in AML cells during respiratory inhibition (**Fig. 2F**). Although respiratory inhibitors have demonstrated promising pre-clinical efficacy when used in combination with venetoclax^52,53^, their clinical success may be undermined by AML cells that can sustain polarization via ATP consumption. In support of this, living MOLM-13_WT_ and MV411_WT_ cells exposed to a combination of venetoclax and respiratory inhibitors (i.e., either IACS and antimycin A, or rotenone and antimycin A) sustained >50% of their basal polarization (**Fig. 4F-G**) (**Supp. Fig. 4E**). Marked inhibition of the remaining polarization was triggered when oligomycin was included, indicating that F_1_-ATPase activity sustains polarization during exposure to a combination of venetoclax and respiratory inhibitors (**Fig. 4F-G**) (**Supp. Fig. 4E**). Based on these data, we hypothesized that oligomycin in combination with either venetoclax or respiratory inhibitors would accelerate AML cell death. To test this, HL60_WT_ and MOLM-13_WT_ cells were treated with combinations of venetoclax, oligomycin, and IACS. When oligomycin and IACS were used in combination, enhanced efficacy was achieved regardless of the presence of venetoclax (**Fig. 4H**); this was consistent with near-complete depolarization of AML cells in response to IACS and oligomycin (**Supp. Fig. 2C**). The addition of IACS with venetoclax also enhanced cytotoxicity in both HL60_WT_ and MOLM-13_WT_ cells (**Fig. 4H-I**), however, a portion of those cells survive by sustaining polarization via the F_1_-ATPase (**Fig. 4F-G**). In the presence of oligomycin, venetoclax reduced the viability of those cells to a larger magnitude (**Fig. 4H-I**), being consistent with sustained polarization by F_1_-ATPase activity following venetoclax exposure (**Fig. 4E**). Collectively, these data suggest that the reversal of the ATP synthase reaction provides the foundation for venetoclax resistance (**Fig. 4J**).

### Chemoresistant AML mitochondria present with a phenotypic shift towards enhanced ATP consumption

Sustaining polarization via F_1_-ATPase activity appears to be an adaptation that opposes the intrinsic pathway of apoptosis, conferring survival to AML cells with reduced respiratory function regardless of the respiratory insult (e.g., by intrinsic complex IV lesions, ETS inhibitors, or venetoclax exposure) (**Fig. 3J**) (**Fig. 4H-I**). Given that venetoclax collapses mitochondrial respiration, we hypothesized that continuous exposure to the drug would positively select for chemoresistant AML subclones that are more reliant on F_1_-ATPase activity to polarize the inner membrane (**Fig. 4C**). To test our hypothesis, we developed in-vitro models of venetoclax resistance by culturing treatment naïve AML cells in venetoclax until they were refractory to the drug (MV411_Vclax_, OCI_Vclax,_ and MOLM-13_Vclax_) (**Fig. 5A**). After multiple rounds of venetoclax selection in culture, a population of AML cells evolved that resisted apoptosis in response to venetoclax (**Fig. 5B**). Having established our in-vitro models of venetoclax resistance, we then evaluated the impact of venetoclax resistance on proteome composition. The abundance of the proton-pumping respiratory complexes (complexes I/III/IV) were downregulated in venetoclax resistant AML cells, an adaptation expected to be deleterious to OxPhos function (**Fig. 5C**). However, venetoclax resistant cells upregulated the abundance of the F_1_-ATPase/ATP synthase (complex V), suggesting an increase in demand for the total amount of F_1_-ATPase proton pumps (**Fig. 5C**). In agreement with this, isolated MV411_Vclax_ mitochondria downregulated the expression of *ATP5IF1*, a specific inhibitor^54–56^ of the F_1_-ATPase (**Fig. 5D**).

**Figure 5.**
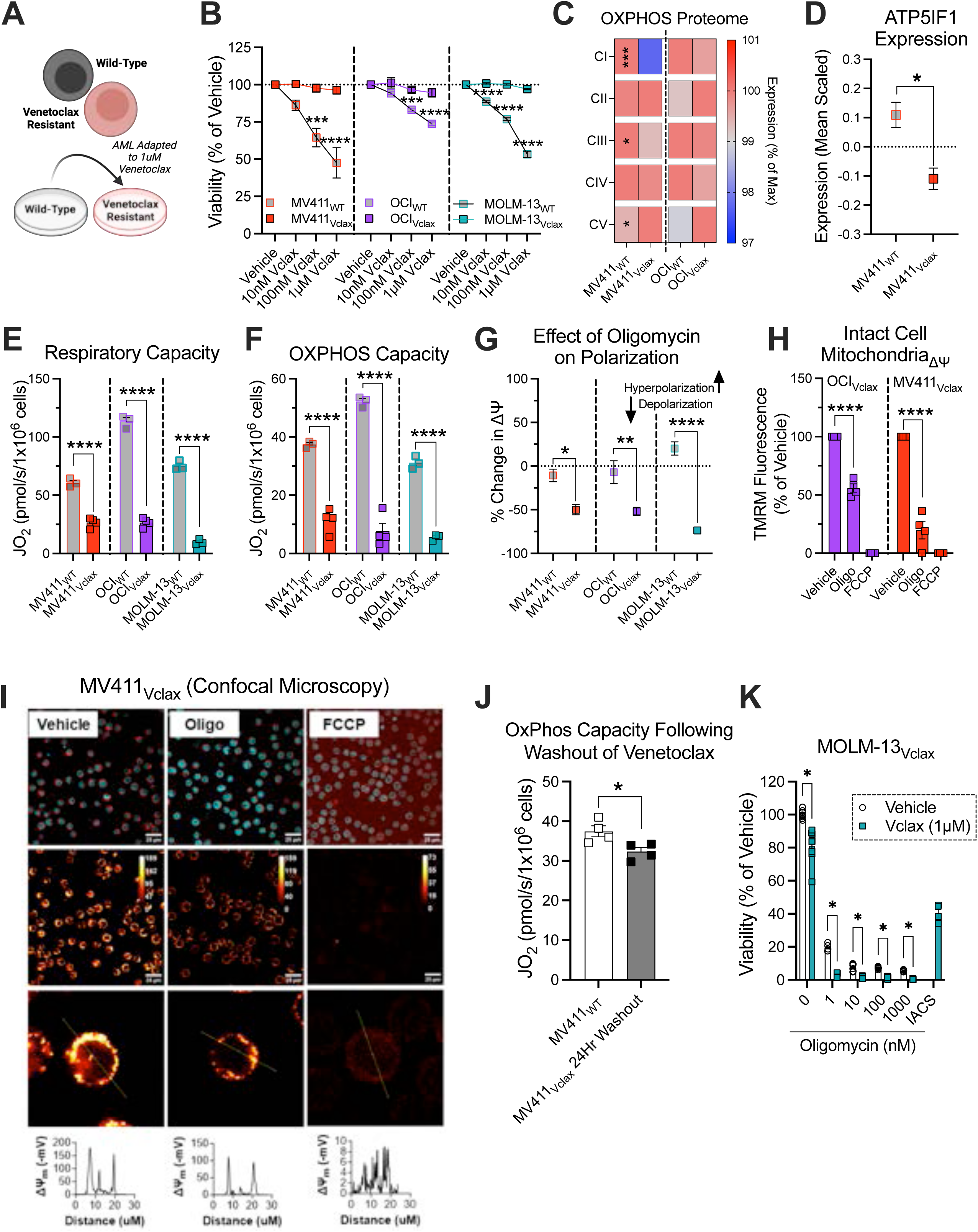
Chemoresistant AML mitochondria present with a phenotypic shift towards enhanced ATP consumption. All experiments were performed using whole intact cells, permeabilized cells, or isolated mitochondria. (A) Schematic depicting generation of venetoclax resistant AML cell lines. (B) Effect of venetoclax on treatment naïve AML and chemoresistant AML cell viability. Viability was measured using propidium iodide (*n* = 3-6 replicates). (C) Comparison of OxPhos proteome in permeabilized treatment naïve AML cells and venetoclax resistant AML cells (*n* = 3 replicates). (D) Comparison of *ATP5IF1* expression in mitochondria isolated from MV411_WT_ and MV411_Vclax_ cells (*n* = 3 replicates). (E) Comparison of respiratory capacity in permeabilized treatment naïve AML cells and venetoclax resistant AML cells (*n* = 3-4 replicates). (F) Comparison of OxPhos capacity in permeabilized treatment naïve AML cells and venetoclax resistant AML cells (*n* = 3-4 replicates). (G) Comparison of oligomycin-induced depolarization in permeabilized treatment naïve AML cells relative to permeabilized venetoclax resistant AML cells (*n* = 3-6 replicates). (H) Flow cytometric analysis of intact cell ΔΨ_m_ in venetoclax resistant AML cells (*n* = 4 replicates). (I) Confocal microscopy images and analysis of MV411_Vclax_ cells in the presence or absence of oligomycin. (J) Comparison of OxPhos capacity in treatment naïve MV411_WT_ cells and MV411_Vclax_ cells after venetoclax has been removed from culture media for 24 hours (*n* = 4 replicates). (K) Effect of IACS and venetoclax, or oligomycin in the presence or absence of venetoclax on MOLM-13_Vclax_ cell viability. Viability was measured using CellTiter-Glo (*n* = 4-8 replicates). Data are presented as mean ±SEM and analyzed by two-way ANOVA (B,I), one-way ANOVA (E,F,G), or unpaired t-test (C,D,H,J,K). *p<0.05, **p<0.01, ***p<0.001, ****p<0.0001.

To determine the impact of venetoclax resistance on the mitochondrial phenotype of AML cells, we compared respiratory capacity and OxPhos capacity of permeabilized treatment naive AML cells and venetoclax resistant AML cells. Venetoclax resistant AML cells presented with a reduction in respiratory capacity and OxPhos capacity, consistent with downregulations in complexes I/III/IV (**Fig. 5C**) (**Fig. 5E-F**) (**Supp. Fig. 5A-F**). Interestingly, cytochrome C addition increased State 4 respiration in MOLM-13_Vclax_ cells, suggesting that venetoclax resistant AML cells can resist apoptosis in response to venetoclax despite incurring damage to the outer mitochondrial membrane (**Supp. Fig. 5C**) (**Supp. Fig. 5F**). We then compared ΔΨ_m_ of permeabilized venetoclax resistant AML cells and treatment naïve AML cells as described previously. Polarization induced by pyruvate/malate was lower in venetoclax resistant AML cells, and this tracked with a greater magnitude of depolarization by oligomycin (**Fig. 5G**) (**Supp. Fig. 5G-I**). To confirm that living venetoclax resistant cells also sustain polarization via F_1_-ATPase activity, intact cell ΔΨ_m_ was assessed. Oligomycin triggered depolarization in living venetoclax resistant AML cells, indicating that F_1_-ATPase activity sustains polarization (**Fig. 5H-I**) (**Supp. Fig. 5J**). Taken together, these data are consistent with our hypothesis that polarization via F_1_-ATPase activity is positively selected for in chemoresistant AML subclones. Given that venetoclax collapses mitochondrial respiration in treatment naïve AML cells, we wanted to determine if venetoclax also collapses mitochondrial respiration in venetoclax resistant AML cells. To test this, venetoclax was removed from MV411_Vclax_ culture media for 24 hours. Following this, OxPhos capacity of MV411_Vclax_ cells was quantified and compared to that of MV411_WT_ cells. Although OxPhos capacity of MV411_WT_ cells remained higher than that of MV411_Vclax_ cells, the removal of venetoclax for 24 hours increased the OxPhos capacity of MV411_Vclax_ cells nearly threefold (**Fig. 5J**) (**Fig. 5F**) (**Supp. Fig. 5K**). Thus, even in chemoresistant cells, venetoclax retains its bioenergetic impact. We hypothesized that venetoclax resistant AML cells can thus be re-sensitized to venetoclax by inhibiting F_1_-ATPase activity. To test this, the effect of oligomycin on MOLM- 13_Vclax_ cell viability was measured in the presence or absence of venetoclax. Venetoclax in combination with oligomycin enhanced cell death of MOLM-13_Vclax_ cells (**Fig. 5K**), consistent with restored sensitivity to venetoclax. Collectively, these data demonstrate that chemoresistant AML mitochondria present with a phenotypic shift towards enhanced ATP consumption, an actionable bioenergetic mechanism critical for cells to evade venetoclax.

### Knockdown of *ATP5IF1* to enhance matrix ATP consumption confers chemoresistance

Continuous exposure to the mitochondrial damaging chemotherapeutic venetoclax positively selected for chemoresistant AML subclones with the following characteristics: 1) upregulated expression of the mitochondrial F_1_-ATPase; 2) downregulation of *ATP5IF1*. Given that *ATP5IF1* specifically inhibits ATP hydrolysis by the F_1_-ATPase^54–56^, we hypothesized that knockdown of *ATP5IF1* would confer resistance to venetoclax. To test this, we used lentiviral particles expressing shRNA targeted to *ATP5IF1* to generate *ATP5IF1* knockdown AML cells lines (“OCI_shIF1_” and “MV411_shIF1_”). Reduced expression of *ATP5IF1* in MV411_shIF1_, relative to MV411_shCtrl_, was confirmed via western blot (**Fig. 6A**). To confirm enhanced F_1_-ATPase activity in *ATP5IF1* knockdown cells, we assessed the sensitivity of uncoupled respiration to inhibition by ΔG_ATP_ as described previously, but without oligomycin. Compared to shRNA control cells, uncoupled respiration in *ATP5IF1* knockdown cells was less sensitive to inhibition during the ΔG_ATP_ titration (**Fig. 6B**), consistent with enhanced F_1_-ATPase activity. When ATPase activity was directly measured in mitochondrial lysates derived from MV411_shIF1_ cells, ATPase activity was enhanced in MV411_shIF1_ cells (**Supp. Fig. 6A**). Having validated our *ATP5IF1* knockdown model, we wanted to determine the impact of *ATP5IF1* knockdown on AML intact cell respiration, respiratory capacity, OxPhos capacity, and fractional OxPhos. *ATP5IF1* knockdown did not result in overt differences in intact cell respiration, respiratory capacity, nor OxPhos capacity (**Fig. 6C-E**) (**Supp. Fig. 6B-C**). However, fractional OxPhos was increased in OCI_shIF1_ cells (**Fig. 6F**). To confirm that F_1_-ATPase activity contributes to polarization in living *ATP5IF1* knockdown AML cells, intact cell ΔΨ_m_ was assessed. Oligomycin triggered depolarization in MV411_shIF1_, indicating that living *ATP5IF1* knockdown cells consume ATP (**Fig. 6G**). After characterizing the mitochondrial phenotype of *ATP5IF1* knockdown AML cells, we tested the hypothesis that *ATP5IF1* knockdown would confer chemoresistance in AML cells. To test this, we measured the impact of *ATP5IF1* knockdown on AML cell viability and colony formation in response to venetoclax. Relative to scrambled shRNA control cells, *ATP5IF1* knockdown cells opposed reductions in cell viability and colony formation in response to venetoclax (**Fig. 6H-J**). Collectively, these data demonstrate that knockdown of *ATP5IF1* boosts matrix F1-ATPase activity, sufficient to confer resistance to venetoclax.

**Figure 6.**
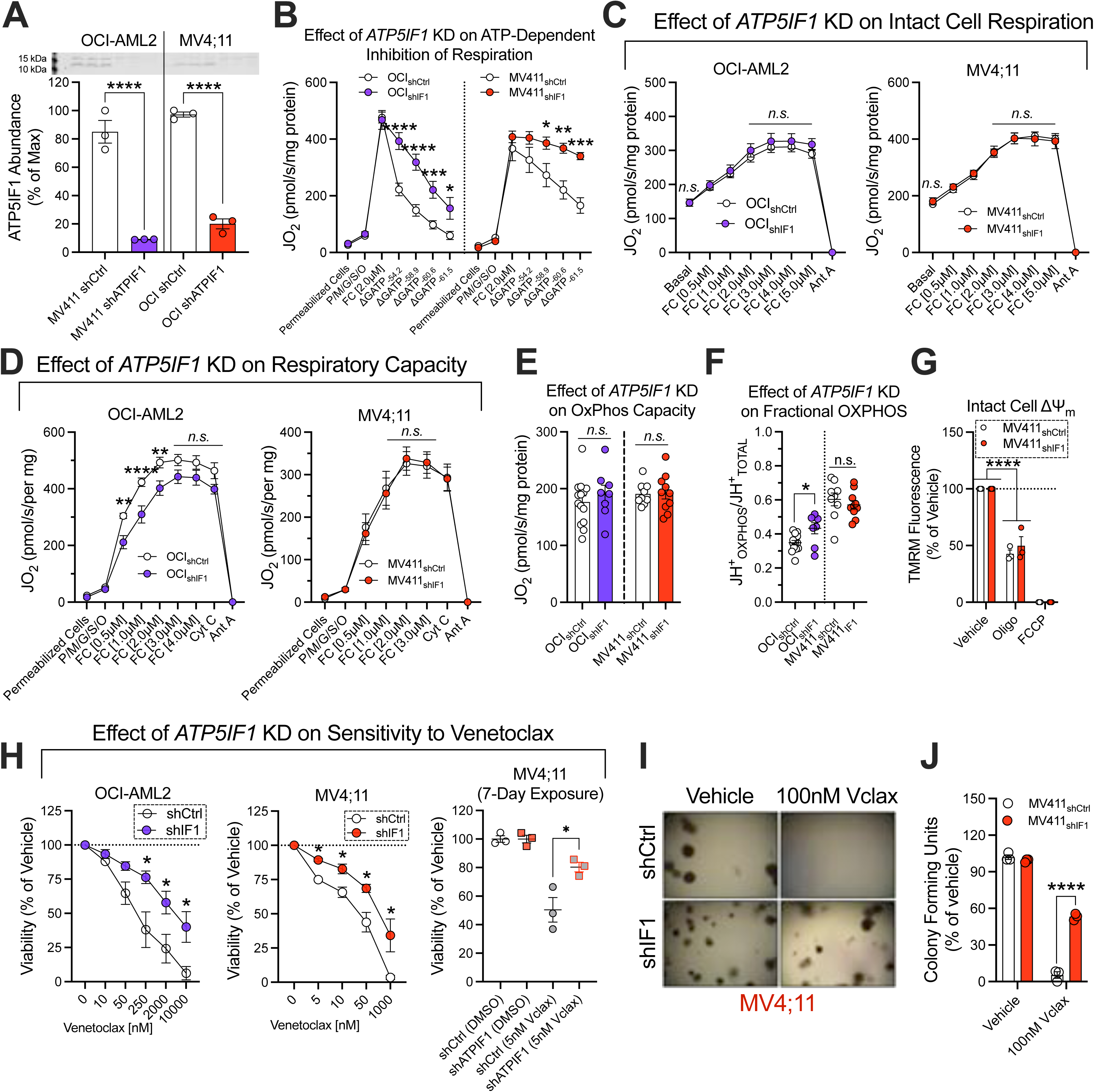
Knockdown of *ATP5IF1* confers chemoresistance. All experiments were performed using whole intact cells, digitonin-permeabilized cells, or isolated mitochondria. (A) Expression of *ATP5IF1* in isolated mitochondria derived from scrambled shRNA control AML cells or *ATP5IF1* knockdown AML cells (*n* = 3 replicates). (B) Impact of *ATP5IF1* knockdown on the inhibition of respiration by ΔG_ATP_ in permeabilized scrambled shRNA control AML cells and permeabilized *ATP5IF1* knockdown AML cells (*n* = 4-8 replicates). (C) Impact of *ATP5IF1* knockdown on intact cell respiration of scrambled shRNA control AML cells and *ATP5IF1* knockdown AML cells (*n* = 8-15 replicates). (D) Impact of *ATP5IF1* knockdown on respiratory capacity in permeabilized scrambled shRNA control AML cells and permeabilized *ATP5IF1* knockdown AML cells (*n* = 8-13 replicates). (E) Impact of *ATP5IF1* knockdown on OxPhos capacity in permeabilized scrambled shRNA control AML cells and permeabilized *ATP5IF1* knockdown AML cells (*n* = 8-14 replicates). (F) Impact of *ATP5IF1* knockdown on Fractional OxPhos in permeabilized scrambled shRNA control AML cells and permeabilized *ATP5IF1* knockdown AML cells (*n* = 8-14 replicates). (G) Flow cytometric analysis of intact cell ΔΨ_m_ in MV411_shCtrl_ and MV411_shIF1_ cells (*n* = 3 replicates). (H) Effect of increasing doses of venetoclax on viability of scrambled shRNA control AML cells and *ATP5IF1* knockdown AML cells after 48 hours, or the effect of venetoclax on MV411_shCtrl_ and MV411_shIF1_ viability after 7 days of exposure. Viability measured using trypan blue viable cell count (*n* = 4 replicates). (I) Effect of venetoclax on colony formation of MV411_shCtrl_ cells and MV411_shIF1_ cells. Colony formation was quantified using CellTiter-Glo (*n* = 3 replicates). (J) Representative image of colony formation of MV411_shCtrl_ cells and MV411_shIF1_ cells in the presence or absence of venetoclax. Data are presented as mean ±SEM and analyzed by two-way ANOVA (B,C,D,G,H,I), one-way ANOVA (A,E), or unpaired t-test (F,H). *p<0.05, **p<0.01, ***p<0.001, ****p<0.0001.

### Overexpression of *ATP5IF1* blunts matrix ATP consumption and enhances chemosensitivity

We hypothesized that overexpression of *ATP5IF1* would restore venetoclax chemosensitivity by inhibiting F_1_- ATPase activity. To test, we used lentiviral particles containing an *ATP5IF1* overexpression construct to generate *ATP5IF1* overexpressing AML cells lines (“OCI_IF1_” and “MV411_IF1_”), in addition to lentivirus control cell lines (“OCI_Ctrl_” and “MV411_Ctrl_”) that overexpress the *Cox8a* mitochondrial targeting sequence tagged to a zsGreen1 fluorescent protein (*Cox8a*MTS:zsGreen1). Following the establishment of stable cell lines, western blotting was used to validate *ATP5IF1* expression in *ATP5IF1* overexpressing AML cells. In OCI_IF1_ cells, *ATP5IF1* expression was increased twofold however, MV411_IF1_ cells did not overexpress *ATP5IF1* (**Fig. 7A-B**). Given that F_1_-ATPase activity is important for AML survival, we hypothesized that *ATP5IF1* expression is selected against in AML cells, similar to that in respiration deficient p^0^ cells^57^. Because p^0^ cells survive by sustaining polarization by consuming ATP^27^, prior work demonstrated that p^0^ cells that survived infection with an *ATP5IF1* overexpression virus were those p^0^ cells that could repress *ATP5IF1* expression^57^. To test our hypothesis, we wanted to determine if non-AML cells would overexpress *ATP5IF1* using the same lentivirus as that used for the overexpression of *ATP5IF1* in AML cells. To do this, we overexpressed *ATP5IF1* in colorectal cancer HCT116 cells (HCT116_IF1_) in addition to lentivirus control HCT116 cells (HCT116_Ctrl_) overexpressing the *Cox8a*-MTS:zsGreen1. Expression of *ATP5IF1* in HCT_IF1_ cells increased nearly fourfold, greater than that in all AML cells lines (**Fig. 7C**) (**Supp. Fig. 6D**) (**Fig. 7D**). These data suggest *ATP5IF1* expression is selected against in AML, consistent with F_1_-ATPase activity being important for AML cell survival.

**Figure 7.**
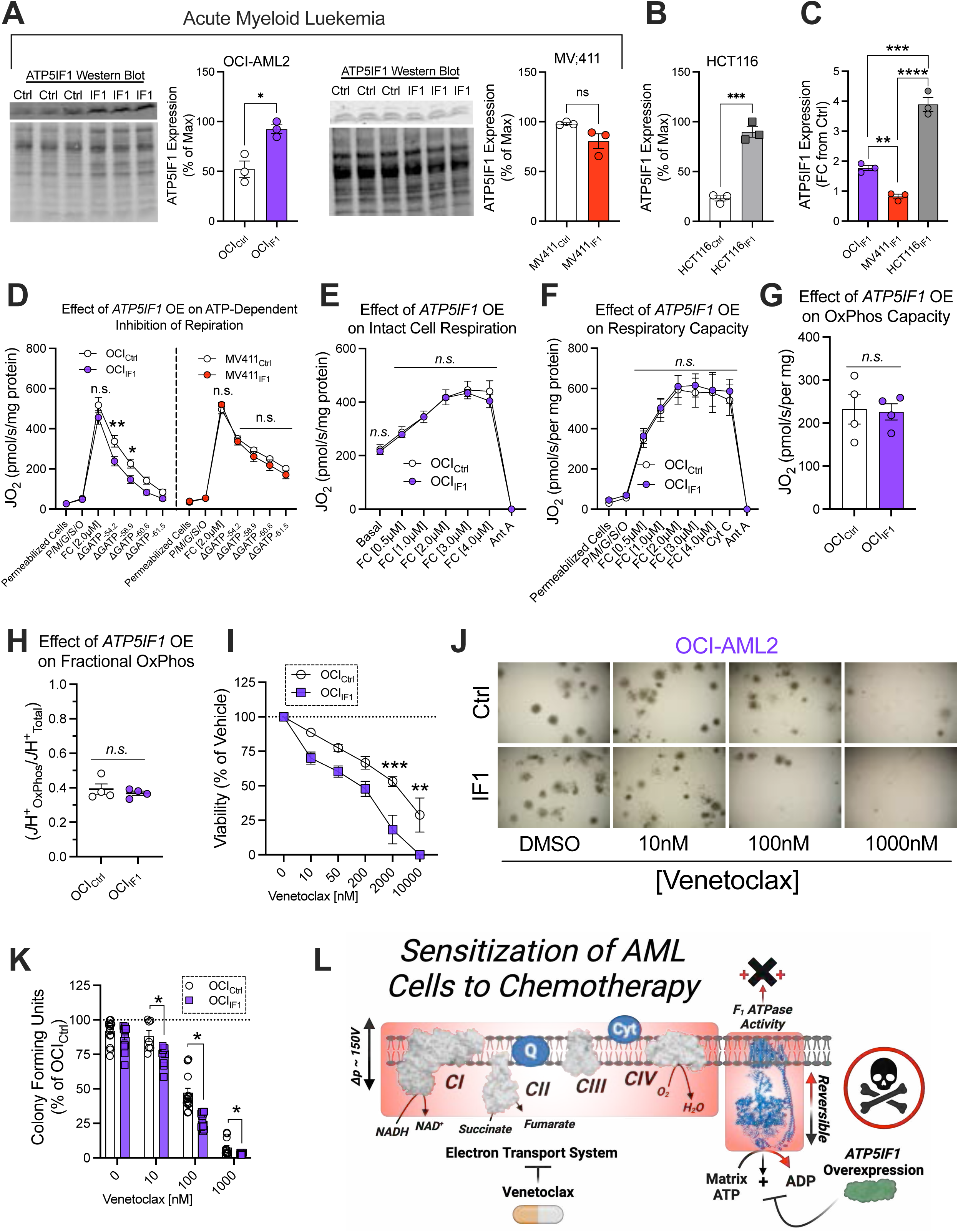
Overexpression of *ATP5IF1* enhances chemosensitivity. All experiments were performed using whole intact cells, digitonin-permeabilized cells, or isolated mitochondria. (A) Expression of *ATP5IF1* in isolated mitochondria derived from lentivirus control AML cells and *ATP5IF1* overexpressing AML cells (*n* = 3 replicates). (B) Expression of *ATP5IF1* in isolated mitochondria derived from lentivirus control HCT116 cells and *ATP5IF1* overexpressing HCT116 cells (*n* = 3 replicates). (C) Overexpression of *ATP5IF1* in isolated mitochondria derived from AML and HCT116 cells infected with *ATP5IF1* overexpression lentivirus represented as fold change from isolated mitochondria derived from lentivirus control AML and HCT116 cells (*n* = 3 replicates). (D) Impact of *ATP5IF1* overexpression on the inhibition of respiration by ΔG_ATP_ in permeabilized lentivirus control AML cells and permeabilized *ATP5IF1* overexpressing AML cells (*n* = 3-13 replicates). (E) Impact of *ATP5IF1* overexpression on intact cell respiration in OCI_Ctrl_ cells and OCI_IF1_ cells (*n* = 5 replicates). (F) Impact of *ATP5IF1* overexpression on respiratory capacity in permeabilized OCI_Ctrl_ cells and permeabilized OCI_IF1_ cells (*n* = 4 replicates). (G) Impact of *ATP5IF1* overexpression on OxPhos capacity in permeabilized OCI_Ctrl_ cells and permeabilized OCI_IF1_ cells. (H) Impact of *ATP5IF1* overexpression on fractional OxPhos in permeabilized OCI_Ctrl_ cells and permeabilized OCI_IF1_ cells (*n* = 4 replicates). (I) Effect of venetoclax on viability of OCI_Ctrl_ cells and OCI_IF1_ cells. Viability measured using trypan blue viable cell count (*n* = 5 replicates). (J) Effect of venetoclax on colony formation of OCI_Ctrl_ cells and OCI_IF1_ cells. Colony formation measured using CellTiter-Glo (*n* = 7-16 replicates). (K) Representative image of colony formation of OCI_Ctrl_ cells and OCI_IF1_ cells in the presence or absence of venetoclax. (L) Schematic depicting mechanism of death of *ATP5IF1* overexpressing AML cells in response to venetoclax. Data are presented as mean ±SEM and analyzed by two-way ANOVA (D,E,F,I), one-way ANOVA (C), or unpaired t-test (A,B,G,H,K) (. *p<0.05, **p<0.01, ***p<0.001, ****p<0.0001.

To confirm that F_1_-ATPase activity is inhibited following *ATP5IF1* overexpression, the sensitivity of uncoupled respiration to inhibition by ΔG_ATP_ was assessed in permeabilized *ATP5IF1* overexpressing cells as described previously. Compared to OCI_Ctrl_ cells, respiration of OCI_IF_ cells was more inhibited by ATP during the ΔG_ATP_ titration (**Fig 7E**). However, respiration of MV411_Ctrl_ and MV411_IF1_ cells were inhibited by ATP at a similar magnitude during the ΔG_ATP_ titration, consistent with no notable increase in *ATP5IF1* expression of MV411_IF1_ cells (**Fig. 7E**). These data suggest that *ATP5IF1* overexpression results in a decrease in F_1_-ATPase activity. When ATPase activity was directly measured in mitochondrial lysates derived from OCI_IF1_ cells, as described previously^44^, ATPase activity was reduced in OCI_IF1_ cells (**Supp. Fig. 6E**). Next, we tested the impact of *ATP5IF1* overexpression on OCI-AML2 intact cell respiration, respiratory capacity, OxPhos capacity, and fractional OxPhos. *ATP5IF1* overexpressing cells did not present with overt differences in intact cell respiration, respiratory capacity, OxPhos capacity, nor fractional OxPhos (**Fig. 7E-H**) (**Supp. Fig. 6F**). Given that neither OxPhos capacity nor fractional OxPhos were overtly inhibited by *ATP5IF1* overexpression, the data are consistent with *ATP5IF1* specifically inhibiting ATP hydrolysis by the F_1_-ATPase^54–56^. We then tested the hypothesis that *ATP5IF1* overexpression would sensitize AML cells to venetoclax. To test, we measured the impact of *ATP5IF1* overexpression on AML cell viability and colony formation in response to venetoclax. Relative to OCI_Ctrl_ cells, OCI_IF1_ cells were more sensitive to reductions in cell viability and colony formation in response to venetoclax (**Fig. 7I-K**). Collectively, these data demonstrate that overexpression of *ATP5IF1* in AML cells enhances sensitivity to venetoclax (**Fig. 7L**).

## DISCUSSION

A diagnosis of AML is abysmal, as over 70% of patients suffering from the disease will succumb to relapse^58,59^. Relapse in AML is caused by the inability of current treatments to eradicate drug resistant subclones which evade chemotherapy, fester below minimal residual disease cutoffs, and ultimately emerge to initiate fatal AML relapse^60^. Therefore, improving AML patient outcomes depends on defining and ultimately thwarting the cellular mechanism(s) that drive chemotherapy resistance. Our prior research linked OxPhos deficiency and AML disease dissemination^22,23^. The current investigation sought to 1) investigate bioenergetic mechanisms that underlie OxPhos deficiency, 2) determine the link between OxPhos deficiency and AML chemoresistant disease, and 3) identify AML-specific mitochondrial drug targets actionable in the context of chemoresistance.

Impaired AML OxPhos flux was caused by pronounced lesions in the oxygen-consuming complex IV of the ETS. Complex IV lesions were masked by increased mitochondrial content in AML cells ^6,22^. Only when respiratory capacity was normalized for mitochondrial content was it clear that maximal respiration intrinsic to each AML mitochondrial unit was lower. Lower respiration resulting from Complex IV lesions would be expected to hamper the ability of the ETS to generate a ΔΨ_m_ sufficient to fuel ATP synthesis. Consistent with this, when OxPhos was assessed in living AML cells, non-canonical ATP hydrolysis by the F_1_-ATPase contributed to polarization under basal conditions. Thus, although living AML cells do respire, they appeared to simultaneously operate the ATP synthase reaction in reverse. The consumption of ATP by the AML F_1_-ATPase was in part related to an intrinsic mechanism where matrix ATP directly disrupts ETS flux, perhaps via phosphorylation or allosteric regulation of matrix enzymes ^61–64^. ATP-dependent inhibition of ETS flux, combined with Complex IV lesions, weakened the voltage potential across the AML inner membrane. Because ΔΨ_m_ drives matrix to cytosol ATP transport via ANT, a weakened voltage potential made AML cells inherently reliant on F_1_-ATPase activity to 1) sustain polarization, and 2) facilitate the clearance of matrix ATP. When we attempted to identify the source of the ATP being consumed by the F_1_-ATPase, inhibition of neither glycolysis by 2-DG, glutaminolysis by CB-839, nor inhibition of ANT by bongkrekic acid were sufficient to fully rescue depolarization in response to oligomycin. Some of the ATP fueling F_1_-ATPase activity may thus be derived from synthesis by ATP in the matrix. Mitochondria contain many copies of the F_1_-ATPase/ATP synthase, and any of those copies can operate either as an ATP synthase or as an F_1_-ATPase, depending on the prevailing mitochondrial voltage potential^31,65,66^ Consistent with our findings in AML cells, *ATP5IF1* expression is necessary to prevent the futile hydrolysis of ATP by the F_1_-ATPase even when OxPhos is occurring at the same time^65,66^. We propose that the interplay between complex IV lesions, matrix ATP, and F_1_-ATPase activity poise the AML mitochondrion to participate in a futile energy cycle where the direction of the F-ATPase/ATP synthase reaction continually responds to fluctuations in ΔΨ_m_ (i.e., akin to an alternating current circuit). By degrading matrix ATP via F_1_-ATPase activity, AML cells oppose the inhibition of respiration by matrix ATP and simultaneously sustain polarization, thereby thwarting intrinsic apoptotic signaling. In doing so, the survival of AML cells is conferred despite intrinsic lesions in the proton pumping respiratory complexes, and at the expense of the “healthy” communication^67,68^ between cellular ΔG_ATP_ demand and mitochondrial metabolism. Given that oxidative metabolism in AML mitochondria appears to be, at least in part, supported by the degradation of matrix ATP via the F_1_-ATPase, we have termed this unique bioenergetic pathway “dephosphorylative oxidation” (i.e., “DephOx), as opposed to canonical “OxPhos” (**Fig. 8**).

**Figure 8.**
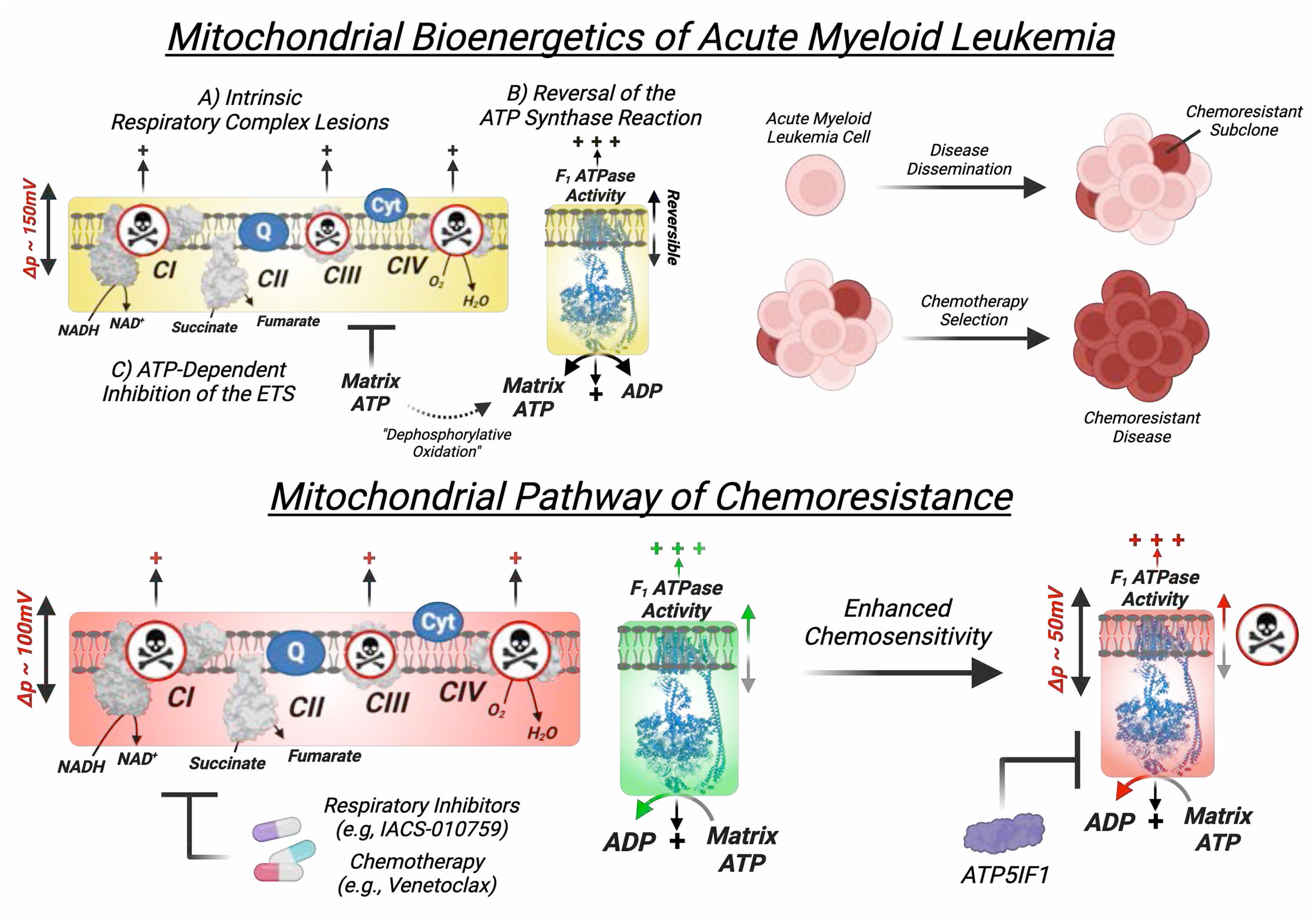
Dephosphorylative Oxidation or, “DephOx”, model of AML mitochondrial bioenergetics. Schematic depicting the Dephosphorylative Oxidation or, “DephOx” model of AML mitochondrial bioenergetics.

The ability of AML mitochondria to both respire and consume ATP via the F_1_-ATPase provides two distinct sources of polarization across the inner membrane, the ETS and F_1_-ATPase. As a result, F_1_-ATPase activity can sustain polarization during respiratory inhibition and thwart apoptotic signaling. In agreement with this, AML cells sustained polarization in response to IACS and antimycin A by consuming ATP via the F_1_- ATPase. Polarization via F_1_-ATPase activity makes AML cells resistant to apoptosis in response to both IACS and antimycin A. Although respiratory inhibitors have demonstrated promising pre-clinical efficacy against AML^52,53,69^ ,the lack of translational efficacy^11,70^ in human trials can be explained, in part, by the inability of current therapies to disrupt polarization originating from the F_1_-ATPase. Related to this, the mitochondrial damaging chemotherapeutic venetoclax is also known to inhibit AML mitochondrial respiration^35,40–41^. AML cells exposed to venetoclax presented with a collapse in mitochondrial respiration that tracked with an inhibition of polarization originating from the ETS. Despite incurring mitochondrial damage and a collapse in mitochondrial respiration, living AML cells exposed to venetoclax remained polarized by consuming ATP via the F_1_-ATPase. Disrupting F_1_-ATPase activity with oligomycin in combination with venetoclax resulted in enhanced toxicity, revealing polarization via the F_1_-ATPase as crucial for AML survival during Bcl-2 inhibition. Consistent with this idea, venetoclax retained its bioenergetic impact on mitochondrial respiration in drug resistant AML. These data suggest that venetoclax should be given to patients continually and in combination with an agent that specifically disrupts polarization by the F_1_-ATPase. Previous reports have demonstrated that AML cell survival is reliant on the activity of the F_1_-ATPase/ATP synthase^71^. Discovered to bind and inhibit ATP synthase at the F_1_ subunit, ammocidin A (i.e., a macrolide) achieved AML cell toxicity in-*vivo* with minimal organ toxicity^71^. The minimal organ toxicity by ammocidin A suggests the exciting possibility that ammocidin A may preferentially inhibit ATP hydrolysis by the F_1_-ATPase, with potential to restore chemosensitivity akin to an *ATP5IF1* mimetic. In our hands, when *ATP5IF1* expression was knocked down in AML cells to enhance F_1_-ATPase activity, resistance to venetoclax was conferred. Conversely, chemosensitivity to venetoclax was enhanced when F_1_- ATPase activity was blunted by overexpression of *ATP5IF1* in OCI-AML2 cells.

Collectively, our data demonstrate that AML cells resist chemotherapy induced apoptosis by hydrolyzing ATP via the F_1_-ATPase to sustain polarization. Polarization of ΔΨ_m_ via F_1_-ATPase activity is generally observed to be a feature of bioenergetically ‘sick’ cells. Given our extensive data revealing AML mitochondria to be bioenergetically ‘sick’, targeting F_1_-ATPase activity in AML cells represents an exciting opportunity to intervene on a unique AML-specific mitochondrial pathway.

## METHODS

All procedures involving human subjects were approved by the Institutional Review Board of the Brody School of Medicine (study ID: UMCIRB 18–001328, UMCIRB 19–002331). All samples were collected in accordance with the Declaration of Helsinki and with informed consent. All procedures on experimental animals were approved by the East Carolina University Institutional Animal Care and Use Committee (Animal Use Protocol Numbers: Q362, Q365).

### Blood collection and isolation of PBMCs

Venous blood from the brachial region of the upper arm was collected from healthy volunteers, between the ages of 18-70 years. Whole blood was collected in sodium-heparinized Cell Preparation Tubes (CPT) (BD Biosciences, Franklin Lakes, NJ) and was then centrifuged at 1,800 x g for 15 min. Mononuclear cells were washed in ammonium-chloride-potassium (ACK) lysis buffer to remove red blood cells and PBMCs were either used immediately, or banked in vapor phase of liquid nitrogen.

### Mononuclear cell isolation from bone marrow aspirates

For primary leukemia samples, bone marrow aspirates were collected from patients undergoing confirmatory diagnosis for a range of hematological malignancies. Patients with confirmed acute myeloid leukemia were enrolled in the study. All patients provided informed consent prior to study enrollment (study ID: UMCIRB 19– 002331). Patient age ranged from 32-78 years. Bone marrow aspirates were collected in sodium-heparinized Cell Preparation Tubes (CPT) (BD Biosciences, Franklin Lakes, NJ) and centrifuged at 1,800 x g for 15 min. Mononuclear cells were isolated and then washed in ACK lysis buffer to remove red blood cells and used immediately for experiments or banked in vapor phase of liquid nitrogen prior to experimentation. For healthy controls, bone marrow aspirates were collected from healthy donors, ages 26-33, supplied by HemaCare (Northridge, CA). Additional heathy bone marrow aspirates were derived from patients enrolled in the study that were discovered to not have a hematological cancer. CD34+ cells were purchased from HemaCare (Northridge, CA).

### Culturing of AML cell lines

MV4-11, HL-60, and KG-1 were originally purchased from ATCC, Manassas, VA. OCI-AML2 cells were a gift from Dr. Mark Minden (Princess Margaret Cancer Center). Mono-Mac-6 (MM6) cells were gifted from Jacqueline Cloos and Carolien van Alphen, VU Medical Center Amsterdam. MOLM-13-YFP-Luc cells (labeled throughout paper as MOLM-13) were generously gifted by Dr. Hong-Gang Wang (Penn State University, Hershey, PA). THP-1 cells were given by Kenichiro Doi and Hong-Gang Wang from Penn State College of Medicine. AML cell lines were cultured in either IMDM (Thermo Fisher Scientific; 31980030) or RPMI-1640 (Thermo Fisher Scientific; 61870-036) supplemented with 10% FBS, and 1% penicillin/streptomycin and incubated at 37°C in 5% CO2. Cells were used for experimentation after reaching an average density 1.0x10^6^ cells/mL, and fresh media was provided 24 hours prior to harvesting.

For experiments involving venetoclax resistance, MV4-11, OCI-AML-2, and MOLM-13 cells were cultured in IMDM (Thermo Fisher Scientific; 31980-030), supplemented with 10% FBS, and 1% penicillin/streptomycin. To model venetoclax resistance, cells were adapted in culture to 1µM venetoclax (Selleck Chemicals; S8048) using methods described previously^74^. AML cells were serially passaged in increasing doses of venetoclax (1nM, 2nM, 5nM, 10nM, 100nM, 1µM). In each passage, AML cells were seeded at a density of 0.5x10^6^ cells/mL and grown to a density of 1x10^6^ cells/mL. After reaching the desired cell density at each passage, AML cells were centrifuged at 300 x *g* for 7 minutes and the culture media was aspirated. AML cells were reseeded at a density of 0.5x10^6^ cells/mL in media supplemented with a higher dose of venetoclax (e.g., from 1nM to 2nM venetoclax). The serial passaging was repeated until AML cells reached a density of 1x10^6^ cell per mL in 1µM venetoclax. Venetoclax resistant AML cells were continually maintained in 1µM venetoclax unless otherwise indicated. Cells were used for experimentation after reaching an average density 1.0x10^6^ cells/mL, and fresh media was provided 24 hours prior to harvesting.

For experiments involving respiration deficient OCI-AML2 p^0^ cells, p^0^ were generated by passaging OCI-AML2 in IMDM (Thermo Fisher Scientific; 31980-030), supplemented with 10% FBS, 1% penicillin/streptomycin, 50ng/mL ethidium bromide, 100 µg/mL pyruvate and 100 µg/mL uridine until a lack of basal respiration was confirmed, similar to that described previously^73^. Cells were used for experimentation after reaching an average density 1.0x10^6^ cells/mL, and fresh media was provided 24 hours prior to harvesting.

### Cell viability

Cells were centrifuged at 300 x *g* for 7 minutes and the culture media was aspirated. Cells were then washed in PBS and centrifuged again at 300 x *g* for 7 minutes. PBS was then aspirated and cells were seeded in either IMDM (Thermo Fisher Scientific; 31980-030) or RPMI-1640 (Thermo Fisher Scientific; 61870-036), supplemented with 10% FBS and 1% penicillin/streptomycin at an average density of 0.2x10^6^ cells/mL. Cell suspension was then loaded into 12 well plates at a volume of 0.5mL and the indicated agents were added to each well relative to a vehicle control. After addition of agents, cells were incubated at 37° C, 5% CO_2,_ for 48 hours. At the end of the 48 hours, cell viability was measured using Trypan Blue (0.4%) (Thermofisher; 15250- 061) viable cell count. Where otherwise indicated, viability was determined by fluorescence measurement using propidium iodide (PI) (Thermo Fisher Scientific; P3566) or luminescence measurement using CellTiter-Glo (Promega; G968A).

For fluorescent measurements of cell viability using propidium iodide, cells were seeded in black-wall, 96-well plates, in growth medium. Water was used as a blank to subtract background fluorescence. After addition of agents (0.2mL final well volume), cells were incubated at 37° C, 5% CO_2,_ for 48 hours. After 48 hours of incubation, positive control cells (used to indicate 100% death) were first permeabilized by addition of 1µl of 10.0 mg/mL digitonin and incubated at 37° C, 5% CO_2,_ for 20 min. Following this, the plates were centrifuged at 1200 x *g* for 10 min. After centrifugation, plates were dumped to remove culture media and 0.1mL of a 5.0µM PI solution in PBS was added to each well. The plate was then incubated at 37° C, 5% CO_2_ for 20 min, and total death (%) was calculated using the ratio of the fluorescence signal of the treatment group and the positive control at 530 nm excitation and 620 nm emission. To determine viability, the total death (%) was subtracted from 100 and normalized relative to the vehicle control. For all viability assays using propidium iodide, each biological replicate was derived from the mean of two technical replicates.

### Colony formation and clonogenicity

Colony formation and clonogenicity of AML cells was performed using Human Methylcellulose Complete Media (bio-techne; 390395). To begin, AML cells were harvested and centrifuged at 300 x *g* for 7 minutes. Following this, AML cells were resuspended at an average density of 1x10^4^ cells/mL to make a 10X stock. The 10X stock of AML cells was then added to separate tubes containing Human Methylcellulose Complete Media to make a 1X cell suspension, and each tube was supplemented with the inhibitors/agents indicated in the figure legends (e.g., venetoclax). Next, 300µL of each cell suspension was seeded in 48-well plate, and any empty wells were filled with sterile, deionized H_2_O to prevent dehydration of the media. The plate was then incubated at 37° C, 5% CO_2_ for 7-10 days. At the end of the incubation period, colony formation was measured by counting colonies using microscopy, or via CellTiter-Glo (Promega; G968A). Representative images were captured using an EVOS FL Auto Imaging System (Thermo Fisher Scientific).

### Quantification and normalization of respiratory flux

Respirometry measurements were conducted using an Oroboros Oxgraph-2k (O2k; Oroboros Instruments, Innsbruck, Austria) in either a 0.5 or 1.0mL reaction volume at 37°C. Data was normalized to viable cell count using Trypan Blue (0.4%) (Thermofisher; 15250-061), unless otherwise indicated. For normalization to total protein, cell suspensions were removed from the O2k chamber following the completion of each assay and placed into microcentrifuge tubes. The cell suspension was then centrifuged at 2,000 x g for 10 min at 4°C. Assay buffer was aspirated and the cell pellet was washed with PBS. Cells were then lysed using low-percentage detergent buffer, CelLytic M (Sigma-Aldrich; C2978), followed by a freeze-thaw cycle on dry ice. The protein concentration was then determined using a Pierce BCA protein assay (Sigma-Aldrich; 71285).

### Intact cell respiration

To assess intact cell respiration, approximately 1-2x10^6^ cells were harvested and centrifuged at 300 x *g*. After centrifugation, cells were washed in PBS and centrifuged again at 300 x *g*. After this, PBS was aspirated and cells were resuspended in either 0.5mL or 1mL Intact Cell Respiratory Media (bicarbonate free IMDM, pH set to 7.4 with HEPES and supplemented with 1% Penicillin/Streptomycin and 10% FBS). Viable cell count was then performed with Trypan Blue (0.4%) (Thermofisher; 15250-061). Basal respiration was then quantified, followed by FCCP titration (FC, 0.5-5µM). Antimycin A (Ant A, 0.5µM) was then added as a negative control to inhibit mitochondrial respiration.

### Permeabilized cell respiratory capacity

To measure respiratory capacity, approximately 1-2x10^6^ cells were harvested and centrifuged at 300 x *g*. After centrifugation, cells were washed in PBS and centrifuged again at 300 x *g*. Cells were then resuspended in either 0.5mL or 1mL of Respiratory Buffer supplemented with creatine (105mM MES potassium salt, 30mM KCl, 8mM NaCl, 1mM EGTA, 10mM KH_2_PO_4_, 5mM MgCl_2_, 0.25% BSA, 5mM creatine monohydrate, pH 7.2). Viable cell count was then performed with Trypan Blue (0.4%) (Thermofisher; 15250-061). Cells were then permeabilized with digitonin [0.02mg/mL]. Respiratory capacity was then assessed by energizing the mitochondria with the indicated carbon substrates - either pyruvate (Pyr or P; 5mM), malate (M; 1mM), octanoyl-carnitine (O; 0.2mM), succinate (Succ or S; 5mM), glutamate (Glut or G; 5mM), or combinations thereof, followed by FCCP titration (FC, 0.5-5µM). Antimycin A (Ant A, 0.5µM) was then added as a negative control to inhibit mitochondrial respiration. Respiratory capacity was determined by the maximal respiration stimulated by any dose of FCCP across the FCCP titration.

### OxPhos respiratory kinetics and OxPhos capacity

OxPhos respiratory kinetics were measured using a modified version of the creatine-kinase clamp (CK Clamp). For experiments using the CK clamp assay, the free energy of ATP hydrolysis (ΔG_ATP_) is calculated using the equilibrium constant for the CK reaction (*K’*_CK_) and is based upon the addition of known concentrations of creatine (CR), phosphocreatine (PCR), and ATP in the presence of excess amounts of CK. Calculation of ΔG_ATP_ at defined PCR concentrations was done using the online resource (https://dmpio.github.io/bioenergetic-calculators/ck_clamp/) as previously described^44^. For all assays using the CK clamp technique, various combinations of carbon substrates and inhibitors were utilized.

To begin to assess OxPhos respiratory kinetics, approximately 1-2x10^6^ cells were harvested and centrifuged at 300 x *g*. After centrifugation, cells were washed in PBS and centrifuged again at 300 x *g*. Cells were then resuspended in either 0.5mL or 1mL of Respiratory Buffer supplemented with creatine (105mM MES potassium salt, 30mM KCl, 8mM NaCl, 1mM EGTA, 10mM KH_2_PO_4_, 5mM MgCl_2_, 0.25% BSA, 5mM creatine monohydrate, pH 7.2). Viable cell count was then performed with Trypan Blue (0.4%) (Thermofisher; 15250- 061). Cells were then permeabilized with digitonin [0.02mg/mL]. OxPhos respiratory kinetics were then assessed by energizing the mitochondria with the indicated carbon substrates - either pyruvate (Pyr or P; 5mM), malate (M; 1mM), octanoyl-carnitine (O; 0.2mM), succinate (Succ or S; 5mM), glutamate (Glut or G; 5mM), or combinations thereof. OxPhos respiratory kinetics were then quantified across a span of physiologically relevant ΔG_ATP_ values (ranging from -54.2 kJ/mol to -61.5 kJ/mol) using the CK clamp (CK [20U/mL], ATP [5mM], PCr [1mM, 6mM, 15mM, 21mM]). At the end of the ΔG_ATP_ titration, oligomycin was added to inhibit ATP synthase, unless otherwise indicated. Following this, maximal respiration was quantified using FCCP uncoupler. At the end of the assay, unless otherwise indicated, antimycin A (Ant A, 0.5µM) was used as a negative control to inhibit mitochondrial respiration. Unless otherwise indicated, the OxPhos capacity of each cell line was calculated as the maximal respiration stimulated by a ΔG_ATP_ value of -54.2 kJ/mol, supported by saturating carbon substrates (P/M/G/S/O). Adaptations of the OxPhos kinetics assay were used to quantify OxPhos capacity supported by either complex I (P/M/G) or complex II (S) linked substrates.

Other substrates and inhibitors used in each assay are indicated in the figure legends: CK [20U/mL], ATP [5mM], PCr [1mM, 6mM, 15mM, 21mM], pyruvate (Pyr or P; 5mM), malate (M; 1mM), octanoyl-carnitine (O; 0.2mM), succinate (Succ or S; 5mM), glutamate (Glut or G; 5mM), oligomycin (Oligo, 0.02µM), FCCP [0.5- 2µM], rotenone (Rot or R, 0.5µM), antimycin A (Ant A, 0.5µM), venetoclax (Vclax, 0.1nM-1µM), FCCP [0.5 µM- 6µM], carboxyatractyloside [5µM], bongkrekic acid [10µg/mL], alamethicin [0.03 mg/mL].

### Isolation of mitochondria from AML cells

Starting material for all isolations was between 300-400 cells. Cells were harvested and centrifuged at 300 x g for 7 min, washed in ice cold PBS, and centrifuged again at 500 x g for 10 min. Cell pellets were then resuspended in Mitochondrial Isolation Buffer with BSA (100mM KCl, 50mM MOPS, 1mM EGTA, 5mM MgSO4, 0.2% BSA, pH 7.1) and homogenized using a borosilicate glass mortar and Teflon pestle. Cell homogenate was centrifuged at 800 x g for 10 min at 4°C. Following this, supernatant was collected and centrifuged at 10,000 x g for 10 min at 4°C to pellet the mitochondrial fraction. The mitochondrial fraction was resuspended in Mitochondrial Isolation Buffer without BSA, transferred to a microcentrifuge tube and centrifuged again at 10,000 x g. The mitochondrial pellet was then resuspended in ∼200µL of Mitochondria Isolation Buffer and protein was quantified using the Pierce BCA assay.

Respiration assays using isolated mitochondria were similar to that described for permeabilized cells. For experiments using permeabilized mitochondria, alamethicin [0.03 mg/mL] was used for permeabilization. In addition to reagents used in the permeabilized cell assays, NADH [4mM] was used to assess complex I-supported respiration in alamethicin-permeabilized mitochondria experiments.

### Isolation of mitochondria from mouse heart

Heart was dissected and placed in ice-cold Mitochondrial Isolation Buffer with BSA (100mM KCl, 50mM MOPS, 1mM EGTA, 5mM MgSO4, 0.2% BSA, pH 7.1). Tissue was then minced, resuspended in 10 ml of Mitochondrial Isolation Buffer and homogenized for ∼10 passes via a Teflon pestle and borosilicate glass vessel. The homogenate was transferred to a 15 ml tube, the volume was brought to 12 ml, and sample was centrifuged at 800 x g for 10 min at 4°C. The supernatant was transferred to a fresh 15mL tube and spun at 10,000 x g for 10 min at 4°C. Mitochondrial pellet was washed in 1.4 ml of Mitochondrial Isolation Buffer without BSA (100mM KCl, 50mM MOPS, 1mM EGTA, 5mM MgSO4, pH 7.1) and spun at 10,000 x g for 10 min at 4°C. The resulting pellet was resuspended in Mitochondrial Isolation Buffer without BSA. Protein content was determined using the Pierce BCA assay.

### Assessment of mitochondrial membrane potential in whole intact cells using flow cytometry

Cells were harvested and resuspended in IMDM (Thermo Fisher Scientific; 31980-030) media at a density of 1x10^6^ cell per mL. After this, cells were treated with either oligomycin [5µM], rotenone [0.5µM], antimycin A [0.5µM], bongkrekik acid [10µg/mL], venetoclax [100nM], CB-839 [1µM], or 2-deoxyglucose [20mM], where indicated, for 15 minutes at 37°C. FCCP [5µM] or alamethicin [0.03 mg/mL] were used as negative controls. The cells were stained with TMRM (10nM) and incubated for 30 minutes at 37°C with 5% CO_2_. Thereafter, DAPI [1µg/mL] was added. Samples were then analyzed with a Cytek Aurora flow cytometer. Data was analyzed by excluding cells determined to be non-viable based on DAPI staining. Given that TMRM is a functional dye, all viable cells were included during the quantification of TMRM fluorescence. FlowJo software v10.9.0 (Treestar, USA) was used for data analysis.

### Assessment of mitochondrial membrane potential in whole intact cells using fluorescence microscopy

Cells were harvested and resuspended in (Thermo Fisher Scientific; 31980-030) media at a density of 1x10^6^ cells/mL. After this, cells were treated with either oligomycin [5µM], rotenone [0.5µM], antimycin A [0.5µM], bongkrekik acid [10µg/mL], or venetoclax [100nM], where indicated, for 15 minutes at 37°C. FCCP [5µM] or alamethicin [0.03 mg/mL] were used as negative controls. The cells were stained with TMRM [100nM] and incubated for 30 minutes at 37°C with 5% CO_2_. Cells were imaged using fluorescence microscopy with an EVOS FL Auto Imaging System (Thermo Fisher Scientific). Quantification of TMRM fluorescence was performed using Cell Profiler^75^ v4.2.5.

For quantification of TMRM fluorescence with Cell Profiler, images taken with fluorescence microscopy were first uploaded into Cell Profiler. Following this, the following modules were added to the pipeline for analyzing each image: 1) IdentifyPrimaryObjects, 2) MeasureObjectIntensity, 3) CalculateMath, 4) ExportToSpreadsheet. The specific parameters for each module (e.g., typical diameter of objects) were defined on a per-experiment basis to optimize the analysis for each image set. In general, cells on the border of the image were excluded from every analysis. The raw analysis was then exported to spreadsheet, and the TMRM fluorescence of each identified object (i.e., each individual cell) was normalized to the average TMRM fluorescence of the vehicle control after subtracting the fluorescence of either FCCP [5µM] or alamethicin [0.03 mg/mL] from all identified objects. Following normalization, the TMRM fluorescence of each identified object in each analyzed image was plotted and represented as individual cells.

For confocal imaging, cells were suspended in IMDM formulation media (Thermo Fisher) containing 50 nM TMRM and 2 μM Hoechst 33342. Cells were plated on glass-bottom dishes (MatTek, Ashland, MA) for imaging. Cells were held in place by pre-coating the glass bottom dishes with Cell Tak (BD, 0.02 mg/mL in 0.1 μM NaHCO3) for 20 minutes. Cells were attached to the plate by centrifugation at 1,000g, without brakes for 5 min. All imaging was performed using an Olympus FV1000 laser scanning confocal microscope (LSCM) with an onstage incubator at 37°C. Acquisition software was Olympus FluoView FSW (V4.2). The objective used was 60X oil immersion (NA = 1.35, Olympus Plan Apochromat UPLSAPO60X(F)). Images were 800 × 800 pixel with 2μs/pixel dwell time, sequential scan mode, resulting in a 4X digital zoom. Hoechst 33342 was excited using the 405 nm line of a multiline argon laser; emission was filtered using a 560 nm dichroic mirror and 420–460 nm barrier filter. TMRM was excited using a 559 nm laser diode; emission was filtered using a 575–675 nm barrier filter. Zero detector offset was used for all images and gain at the detectors was kept the same for all imaging. The pinhole aperture diameter was set to 105 μm (1 Airy disc). Images were analyzed using Fiji^76^. Spatial resolution was measured using sub-resolution fluorescent beads (Thermo Fisher) and curve fitting was performed using the MetroloJ plugin in Fiji. 16-bit images were made into a composite. ΔΨ_m_ was measured using non-quenching concentrations of TMRM. Out-of-field images from the axial dimension were trimmed from each stack. Image stacks were then background-corrected using darkfield images. Median background intensity (I_B_) was measured by inverting a Huang auto threshold for each image stack. The estimated electrical potential difference relative to the median value of the extra-mitochondrial signal was then mapped onto each pixel in each image using a math macro:

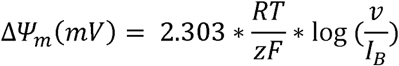

where R is the universal gas constant (J/mol*K), T is the absolute temperature (310.15K), z is the +J1 charge of TMRM, F is Faraday’s constant (C), and v is the gray value of the designated pixel. A Huang auto threshold was then applied to each image in the stack, and median and maximum values for each threshold were obtained. This measurement method reduced biasing of the total measured signal toward low-intensity pixel values.

### Assessment of membrane potential of isolated mitochondria using flow cytometry

Mitochondria isolated from mouse heart or HL60 cells were treated with either 1) H_2_O (Vehicle), 2) pyruvate (Pyr or P; 5mM) and malate (M; 1mM), 3) P/M, CK [20U/mL], ATP (5mM), and PCr [21mM], 4) P/M, CK [20U/mL], ATP [5mM], PCr [21mM] and Oligo [5µM], or 5) P/M, CK [20U/mL], ATP [5mM], PCr [21mM], Oligo [5µM] and FCCP [5µM] for 15 minutes at room temperature. The cells were stained with TMRM [10nM] and incubated for 30 minutes at 37°C with 5% CO_2_. Samples were then analyzed with a Cytek Aurora flow cytometer. TMRM fluorescence intensity was quantified using FlowJo software v10.9.0 (Treestar, USA).

### Assessment of mitochondrial membrane potential in permeabilized cells

Mitochondrial membrane potential in permeabilized cells was performed using methods described previously^44^ with some adaptations. Fluorescent determination of the mitochondrial membrane potential was performed using a QuantaMaster Spectrofluorometer (QM-400; Horiba Scientific). Polarization was determined using TMRM fluorescence operating in quench mode, as described previously^44^. Cells were harvested, centrifuged at 300 x g for 7 minutes, then washed with PBS and centrifuged again at 300 x g. Following this, cells were resuspended in 1mL of Respiratory Buffer supplemented with creatine (105mM MES potassium salt, 30mM KCl, 8mM NaCl, 1mM EGTA, 10mM KH_2_PO_4_, 5mM MgCl_2_, 0.25% BSA, 5mM creatine monohydrate, pH 7.2). Viable cell count was then performed with Trypan Blue (0.4%) (Thermofisher; 15250-061). Following this, cells were permeabilized with digitonin (Digi; 0.02mg/mL) for 10 minutes at room temperature, with gentle rocking. From that permeabilized cell suspension, ∼3 million cells were added to a microcentrifuge tube and centrifuged at 1,500 x g for 10 minutes. The permeabilized cell pellet was resuspended in fresh Respiratory Buffer and loaded with TMRM [200nM]. Optimal excitation and emission parameters for each cell type were determined using an excitation/emission scan to determine the fluorescence ratio, as described previously^44^. Membrane potential was then quantified using a fluorescence ratio of the experimentally determined excitation and emission parameters.

In general, permeabilized cell membrane potential assays were similar to those used in respiration assays. Substrates and inhibitors used in each assay are indicated in the figure legends: CK [20U/mL], ATP [5mM], PCr [1mM, 6mM, 15mM, 21mM], pyruvate (Pyr or P; 5mM), malate (M; 1mM), oligomycin (Oligo, 0.02µM), FCCP [5µM], rotenone (Rot or R, 5-100nM), antimycin A (Ant A, 0.5µM), venetoclax (Vclax, 0.1nM-1µM), FCCP [5µM], carboxyatractyloside [5µM].

### Assessment of ATP-dependent inhibition of respiration and polarization using permeabilized cells

To assess the inhibition of respiration by ATP, cells were harvested and centrifuged at 300 x *g*. After centrifugation, cells were washed in PBS and centrifuged again at 300 x *g*. Cells were then resuspended in either 0.5mL or 1mL of Respiratory Buffer supplemented with creatine (105mM MES potassium salt, 30mM KCl, 8mM NaCl, 1mM EGTA, 10mM KH_2_PO_4_, 5mM MgCl_2_, 0.25% BSA, 5mM creatine monohydrate, pH 7.2). Viable cell count was then performed with Trypan Blue (0.4%) (Thermofisher; 15250-061). Cells were then permeabilized with digitonin [0.02mg/mL]. Following permeabilization, cells were energized with saturating carbon substrates - pyruvate (Pyr or P; 5mM), malate (M; 1mM), octanoyl-carnitine (O; 0.2mM), succinate (Succ or S; 5mM), and glutamate (Glut or G; 5mM). Uncoupled respiration was stimulated with FCCP [1.5- 2µM]. In wild-type AML cells, oligomycin [0.02µM] or no oligomycin was added to inhibit the reversal of ATP synthase. The inhibition of uncoupled respiration by ATP was then quantified across a span of physiologically relevant ΔG_ATP_ values (ranging from -54.2 kJ/mol to -61.5 kJ/mol) using the CK clamp (CK [20U/mL], ATP [5mM], PCr [1mM, 6mM, 15mM, 21mM]). When this assay involved cells that have undergone genetic manipulation of *ATP5IF1* expression, oligomycin was not used.

To assess the inhibition of polarization by ATP, cells were harvested and centrifuged at 300 x *g*. After centrifugation, cells were washed in PBS and centrifuged again at 300 x *g*. Cells were then resuspended in either 0.5mL or 1mL of Respiratory Buffer supplemented with creatine (105mM MES potassium salt, 30mM KCl, 8mM NaCl, 1mM EGTA, 10mM KH_2_PO_4_, 5mM MgCl_2_, 0.25% BSA, 5mM creatine monohydrate, pH 7.2). Viable cell count was then performed with Trypan Blue (0.4%) (Thermofisher; 15250-061). Cells were then permeabilized with digitonin [0.02mg/mL] and gently rocked for 10 minutes. From that permeabilized cell suspension, ∼3 million cells were added to a microcentrifuge tube and centrifuged at 1,500 x g for 10 minutes. The permeabilized cell pellet was resuspended in fresh Respiratory Buffer and loaded with TMRM [200nM] and the membrane potential was assessed, similar to that described in the permeabilized cell membrane potential assay with some modifications. Oligomycin [0.02µM] was added to inhibit ATP synthase and either carboxyatractyloside [5µM] or no carboxyatractyloside was added to inhibit the adenine nucleotide translocase (ANT). Mitochondria were then polarized with pyruvate (Pyr or P; 5mM) and malate (M; 1mM). The inhibition of polarization by ATP was then quantified across a span of physiologically relevant ΔG_ATP_ values (ranging from - 54.2 kJ/mol to -61.5 kJ/mol) using the CK clamp (CK [20U/mL], ATP [5mM], PCr [1mM, 6mM, 15mM, 21mM]). Antimycin A (Ant A, 0.5µM) was then added followed by FCCP [5µM] as a negative control.

### Lentiviral *ATP5IF1* knockdown in AML cells

OCI-AML2 or MV4-11 cells were cultured in IMDM (Thermo Fisher Scientific; 31980-030) supplemented with 10% FBS and 1% penicillin/streptomycin and incubated at 37°C in 5% CO2. Human shRNA lentiviral particles cloned into pLKO.1 vector (18mer *ATP5IF1*-specific shRNA 5’ GGCTGAAGAGGAACGATA 3’), and one scramble control (0.5mL each,>10^7^ TU/mL); were purchased from Addgene (CAT#: 10878). To facilitate infection, AML cells and lentiviral particles were co-cultured at a seeding density of 1x10^6^ cells per mL in a Retronectin dish (Takara Bio; T110A). Following 72 hours of infection, cultures were then subjected to puromycin selection by continuous exposure to puromycin [1-2µg/mL] in the culture media.

To confirm knockdown of *ATP5IF1* expression, mitochondria were first isolated from scrambled shRNA control cells and *ATP5IF1* knockdown AML cells. Mitochondria were then lysed using Pierce RIPA Buffer (Thermo Fisher Scientific; 89900), and were supplemented with 1X Halt Protease and Phosphatase Inhibitor Cocktail, EDTA-free (Thermo Fisher Scientific; 78441). Mitochondrial lysates underwent a standard western blot protocol using a Mini-PROTEAN TGX stain-free gel (Bio-Rad; 4568096) run at 150V. An anti-*ATP5IF1* antibody (Thermo Fisher Scientific; A21355) was used to bind *ATP5IF1* protein on the blot membrane, and a secondary anti-mouse antibody was used for either fluorescent or luminescent detection of *ATP5IF1* expression. Western blot images were analyzed by densitometry using ImageJ v1.53 software, and the density of each *ATP5IF1* band was normalized to the density of total protein in each lane to quantify *ATP5IF1* expression.

### Lentiviral *ATP5IF1* overexpression in AML and HCT116 cells

AML cells were cultured in IMDM (Thermo Fisher Scientific; 31980-030) supplemented with 10% FBS and 1% penicillin/streptomycin, and HCT116 cells were cultured in RPMI-1640 (Thermo Fisher Scientific; 61870-036) supplemented with 10% FBS, and 1% penicillin/streptomycin and incubated at 37°C in 5% CO2. The Pantropic ViraSafe^TM^ Universal Lentiviral Expression System (Cell Biolabs, Inc.; VPK-211-PAN) was used to establish cells overexpressing the human *ATP5IF1* and the *Cox8a*MTS:zsGreen1, with *Cox8a* also being derived from human. zsGreen1 is a human codon-optimized variant of Zoanthus sp. Green fluorescent protein zsGreen. The plasmid used the Ef1-alpha promoter to drive expression of both *ATP5IF1* and *Cox8a*MTS:zsGreen1. For antibiotic selection, a puromycin resistance gene was also expressed in the plasmid, driven by the human phosphoglycerate kinase 1 promoter. To facilitate infection, either AML cells were co-cultured with lentiviral particles at a seeding density of 1x10^6^ cells per mL in a Retronectin dish (Takara Bio; T110A). HCT116 cells were seeded at a density of 0.2x10^6^ cells per mL in a six well plate. Following 72 hours of infection for all cell types, cultures were then subjected to puromycin selection by continuous exposure to puromycin [1-2µg/mL] in the culture media.

To confirm overexpression of *ATP5IF1*, mitochondria were first isolated from *Cox8a*MTS:zsGreen1 control cells and *ATP5IF1* overexpressing cells. Mitochondria were then lysed using Pierce RIPA Buffer (Thermo Fisher Scientific; 89900), and were supplemented with 1X Halt Protease and Phosphatase Inhibitor Cocktail, EDTA-free (Thermo Fisher Scientific; 78441). Mitochondrial lysates underwent a standard western blot protocol using a Mini-PROTEAN TGX stain-free gel (Bio-Rad; 4568096) run at 150V. An anti-*ATP5IF1* antibody (Thermo Fisher Scientific; A21355) was used to bind *ATP5IF1* protein on the blot membrane, and a secondary anti-mouse antibody was used for either fluorescent or luminescent detection of *ATP5IF1* expression. Western blot images were analyzed by densitometry using ImageJ v1.53 software, and the density of each *ATP5IF1* band was normalized to the density of total protein in each lane to quantify *ATP5IF1* expression.

### Determination of ATP hydrolytic activity of Complex V

ATPase activity was assessed as described previously^78^ with slight modifications. Mitochondrial lysates for the assay were prepared via dilution of the final isolated mitochondrial suspensions in CelLytic M (Sigma-Aldrich; C2978) at a protein concentration of 2mg/ml. Buffer for the assay was ATPase Assay Buffer, supplemented with lactate dehydrogenase/pyruvate kinase [10U/mL], phosphoenoyl-pyruvate [5mM], rotenone [0.005mM] and NADH [0.2mM]. Assay buffer (200 μL) was loaded into individual wells of a 96-well plate, followed by mitochondrial lysate (2 μg/well). The assay was initiated with the addition of ATP [5mM]. In the assay, NADH oxidation and ATP hydrolysis occur at a 1:1 stoichiometry and thus ATPase activity (represented as a % of control cells) was determined via tracking the degradation in the NADH auto-fluorescence (Ex:Em 376/450) signal upon ATP addition.

### Determination of intact cell respiration normalized to mitochondrial content

Experimentation on OCI-AML2, MV4-11, and Mobilized PBMC cells was conducted after thawing cells directly from stored, frozen aliquots (-80°C). Each cell type was thawed rapidly in a warm bath (37°C) and then were centrifuged at 300 x *g*. The supernatant was aspirated and the cells were resuspended in PBS. Viable cell count was then performed with Trypan Blue (0.4%) (Thermofisher; 15250-061). From that suspension, 1x10^6^ total cells were placed in microcentrifuge tubes and were stored at (-80°C) for proteomics preparation. The remaining cells were centrifuged again at 300 x *g*. After this, PBS was aspirated and cells were resuspended in either 0.5mL or 1mL Intact Cell Respiratory Media (supplemented with 1% Penicillin/Streptomycin and 10% FBS, pH 7.4). Viable cell count was again performed with Trypan Blue (0.4%) (Thermofisher; 15250-061) for normalization of intact cell respiration. Basal respiration was then quantified, followed by FCCP titration (FC, 0.5-5µM). Antimycin A (Ant A, 0.5µM) was then added as a negative control to inhibit mitochondrial respiration. At the end of each assay, cell suspensions were removed from the O2k chamber and placed into microcentrifuge tubes. The cell suspension was then centrifuged at 2,000 x g for 10 min at 4°C. Assay buffer was aspirated and the cell pellet was washed with PBS. Cells were then lysed using low-percentage detergent buffer, CelLytic M (Sigma-Aldrich; C2978), followed by a freeze-thaw cycle on dry ice. The protein concentration was then determined using a Pierce BCA protein assay (Sigma-Aldrich; 71285) and intact cell respiration was normalized to milligrams of total intact cell protein.

The frozen 1x10^6^ cell aliquots that were stored from each cell type (i.e., OCI-AML2, MV4-11, and Mobilized PBMC cells) were later used for proteomic analysis. Following proteomics analysis, the ratio of the mitochondrial protein to total protein was used to calculate the mitochondrial enrichment factor (MEF) which provides an estimate of mitochondrial content^79^. The MEF was then used to normalize intact cell respiration (after normalization to milligrams of total intact cell protein) to the rate of respiration per milligram of total mitochondrial protein.

### Sample prep for label-free proteomics

Cell pellets or isolated mitochondria pellets were lysed in urea lysis buffer (8M urea in 40mM Tris, 30mM NaCl, 1mM CaCl_2_, 1 x cOmplete ULTRA mini EDTA-free protease inhibitor tablet; pH=8.0), as described previously^80^. The samples were subjected to two freeze-thaw cycles, and sonicated with a probe sonicator in three 5s bursts (Q Sonica #CL-188; amplitude of 30). Samples were centrifuged at 10,000 x g for 10min at 4°C. Protein concentration was determined by BCA. Equal amounts of protein were reduced with 5mM DTT at 37°C for 30min, and then alkylated with 15mM iodoacetamide for 30min in the dark. Unreacted iodoacetamide was quenched with DTT [15mM]. Reduction and alkylation reaction were carried out at room temperature. Initial digestion was performed with Lys C (1:100 w:w) for 4hr at 32°C. Following dilution to 1.5M urea with 40mM Tris (pH=8.0), 30mM NaCl, 1mM CaCl_2_, samples were digested overnight with sequencing grade trypsin (50:1 w/w) at 32°C. Samples were acidified to 0.5% TFA and then centrifuged at 4,000 x g for 10min at 4°C. Supernatant containing soluble peptides was desalted and then eluate was frozen and subjected to speedvac vacuum concentration.

### nLC-MS/MS for label-free proteomics

Final peptides were resuspended in 0.1% formic acid, quantified (Thermo Fisher Scientific; 23275), and diluted to a final concentration of 0.25µg/µL. Samples were subjected to nanoLC-MS/MS analysis using an UltiMate 3000 RSLCnano system (Thermo Fisher Scientific) coupled to a Q Exactive Plus Hybrid Quadrupole-Orbitrap mass spectrometer (Thermo Fisher Scientific) via a nanoelectrospray ionization source. For each injection, 4µL (1µg) of sample was first trapped on an Acclaim PepMap 100 20mm × 0.075mm trapping column (Thermo Fisher Scientific; 164535; 5μL/min at 98/2 v/v water/acetonitrile with 0.1% formic acid). Analytical separation was performed over a 95min gradient (flow rate of 250nL/min) of 4-25% acetonitrile using a 2µm EASY-Spray PepMap RSLC C18 75µm × 250mm column (Thermo Fisher Scientific; ES802A) with a column temperature of 45°C. MS1 was performed at 70,000 resolution, with an AGC target of 3x10^6^ ions and a maximum injection time (IT) of 100ms. MS2 spectra were collected by data-dependent acquisition (DDA) of the top 15 most abundant precursor ions with a charge greater than 1 per MS1 scan, with dynamic exclusion enabled for 20s. Precursor ions isolation window was 1.5m/z and normalized collision energy was 27. MS2 scans were performed at 17,500 resolution, maximum IT of 50ms, and AGC target of 1x10^5^ ions.

### Data analysis for label-free proteomics

As described previously^80^ with some modification, Proteome Discoverer 2.2 (PDv2.2) was used for raw data analysis, with default search parameters including oxidation (15.995 Da on M) as a variable modification and carbamidomethyl (57.021 Da on C) as a fixed modification. Data were searched against the Uniprot Homo Sapiens reference proteome (Proteome ID: UP000005640), as well as the Human Mito Carta 3.0 database^81^. PSMs were filtered to a 1% FDR and grouped to unique peptides while maintaining a 1% FDR at the peptide level. Peptides were grouped to proteins using the rules of strict parsimony and proteins were filtered to 1% FDR. Peptide quantification was done using the MS1 precursor intensity. Imputation was performed via low abundance resampling. Using only high confidence master proteins, mitochondrial enrichment factor (MEF) was determined as previously described^79^ by comparing mitochondrial protein abundance (i.e., proteins identified to be mitochondrial by cross-reference with the MitoCarta 3.0 database) to total protein abundance.

### Statistical evaluation of label-free proteomics

All proteomics samples were normalized to total protein abundance, and the protein tab in the PDv2.2 results was exported as a tab delimited .txt. file and analyzed. Protein abundance was converted to the Log_2_ space. For pairwise comparisons, cell mean, standard deviation, p-value (p; two-tailed Student’s t-test, assuming equal variance), and adjusted p-value (Benjamini Hochberg FDR correction) were calculated. Throughout the paper, differences between groups were assessed by t-test, or one-way ANOVA, followed by Tukey’s test where appropriate using GraphPad Prism 8 software (Version 8.4.2). Other statistical tests used are described in the figure legends. Statistical significance in the figures is indicated as follows: *p < 0.05; **p < 0.01; ***p < 0.001; ****p < 0.0001.

### Statistical analysis and software

Statistical analysis was performed using GraphPad Prism 8.4. All data are represented as mean ± SEM and analysis were conducted with a significance level set at p<0.05. Details of statistical analysis are included within figure legends. Figures were created with BioRender.com and GraphPad Prism 10.1.1.

### Data availability

All data from the manuscript are available upon request. In addition, all data are available in the source data files provided with this paper (**Supplemental Table 1**). All raw data for proteomics experiments is available online using accession number “PXD051129” for Proteome Xchange^82^ and accession number “JPST003019” for jPOST Repository^83^. Proteomics data are also available in **Supplemental Table 2**.

## AKNOWELDGEMENTS

This work was supported by DOD-W81XWH-19-1-0213 (K.H.F.W.) and NIH P01 CA171983 (M.C.C./T.P.L.). In addition, this work was supported by a UNC Lineberger Comprehensive Cancer Center Developmental Award which is supported in part by P30 CA016086 Cancer Center Core Support Grant. During the performance of this work, K.H.F-W. relocated from East Carolina University to Wake Forest University School of Medicine.

---

**Supplementary Figure 1.**
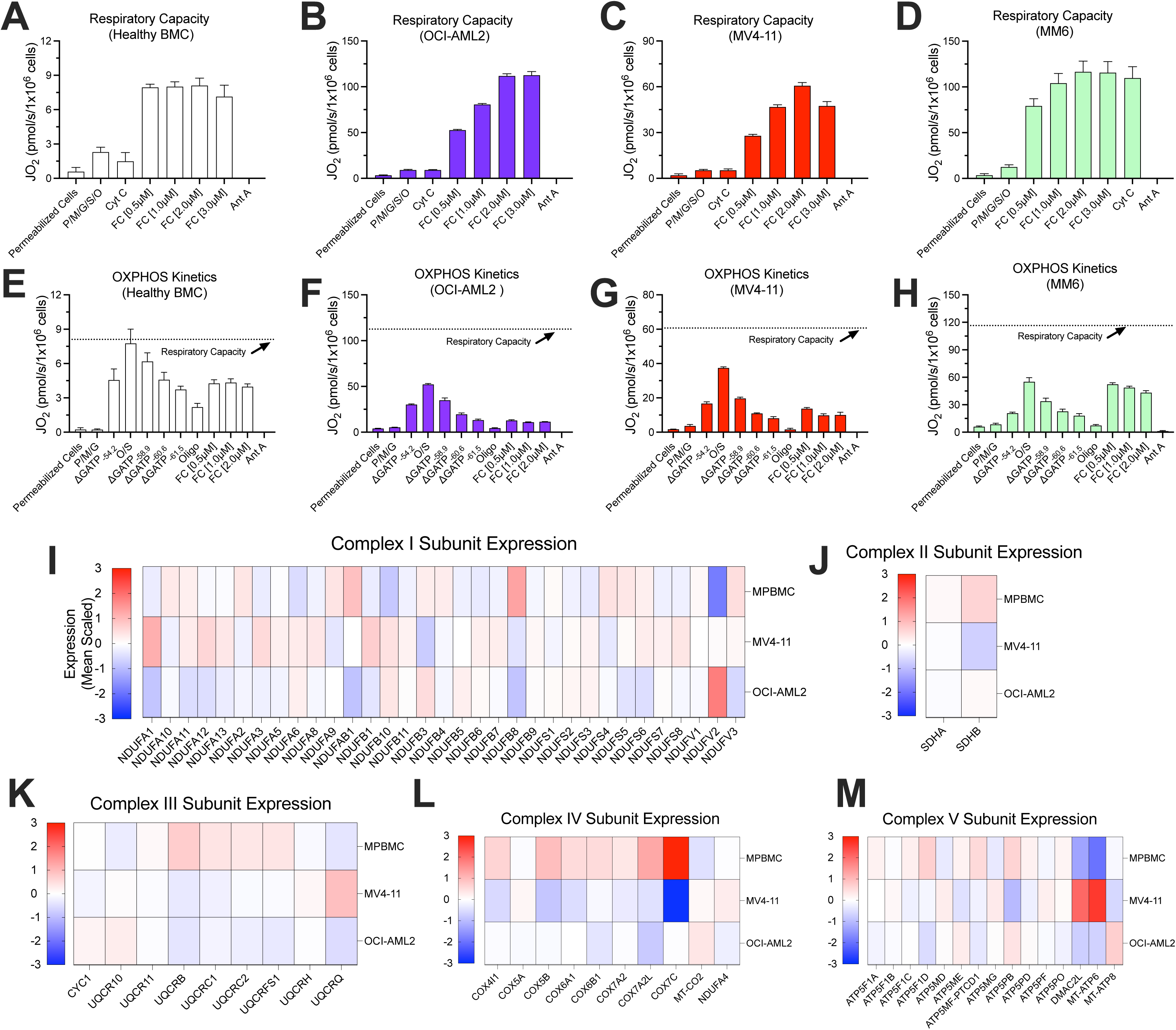
Respiratory capacity and OXPHOS kinetics profiling of AML and healthy hematopoietic cells. (Related to Figure 1) All experiments were performed using whole intact or digitonin-permeabilized cells. Respiratory capacity was measured using (A) Healthy BMC (*n* = 3 replicates), (B) OCI_WT_ (*n* = 3 replicates), (C) MV4-11 (*n* = 3 replicates), (D) MM6 cells. OXPHOS respiratory kinetics were measured using (E) Healthy BMC (*n* = 3 replicates), (F) OCI-AML2 (*n* = 3 replicates), (G) MV4-11 (*n* = 3 replicates), (H) MM6 cells (*n* = 3 replicates). (I) Comparison of Complex I subunit expression in Mobilized PBMC and AML cells (*n* = 3 replicates). (J) Comparison of Complex II subunit expression in Mobilized PBMC and AML cells (*n* = 3 replicates). (K) Comparison of Complex III subunit expression in Mobilized PBMC and AML cells (*n* = 3 replicates). (L) Comparison of Complex IV subunit expression in Mobilized PBMC and AML cells (*n* = 3 replicates). (M) Comparison of Complex V subunit expression in Mobilized PBMC and AML cells (*n* = 3 replicates).

**Supplementary Figure 2.**
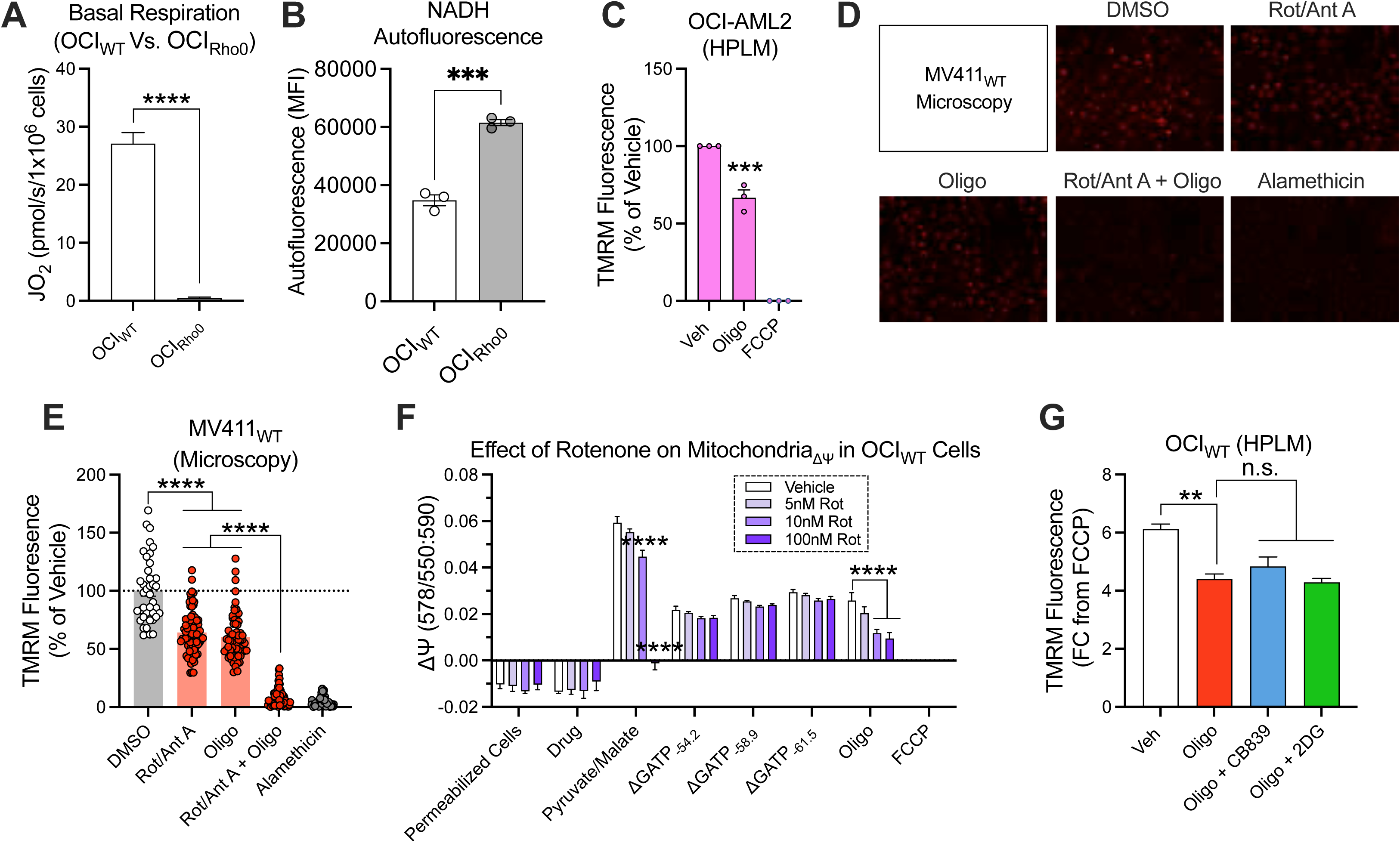
Extended validation of p^0^ OCI-AML2 cells and F_1_-ATPase activity in AML. (Related to Figure 2) All experiments were performed using whole intact cells or isolated mitochondria. (A) Comparison of basal respiration of OCI_Rho0_ or OCI_WT_ cells (*n* = 4 replicates). (B) Comparison of NADH autofluorescence of OCI_Rho0_ or OCI_WT_ cells (*n* = 3 replicates). (C) Flow cytometric analysis of intact cell ΔΨ_m_ in OCI_WT_ cells assayed in human plasma-like media (HPLM) (*n* = 3 replicates). (D) Fluorescent microscopy representative images of intact cell ΔΨ_m_ in MV411_WT_ cells. (E) Fluorescent microscopy analysis of intact cell ΔΨ_m_ in MV411_WT_ cells (*n* = 39-154 cells). (F) Effect of rotenone on permeabilized OCI_WT_ cell ΔΨ_m_ (*n* = 4 replicates). (G) Flow cytometric analysis of intact cell ΔΨ_m_ in OCI_WT_ cells in the presence of oligomycin, or that in combination with 2-DG or CB-839 (*n* = 3 replicates). Data are presented as mean ±SEM and analyzed by two-way ANOVA (E,F,G), one-way ANOVA (C), or unpaired t-test (A,B). *p<0.05, **p<0.01, ***p<0.001, ****p<0.0001.

**Supplementary Figure 3.**
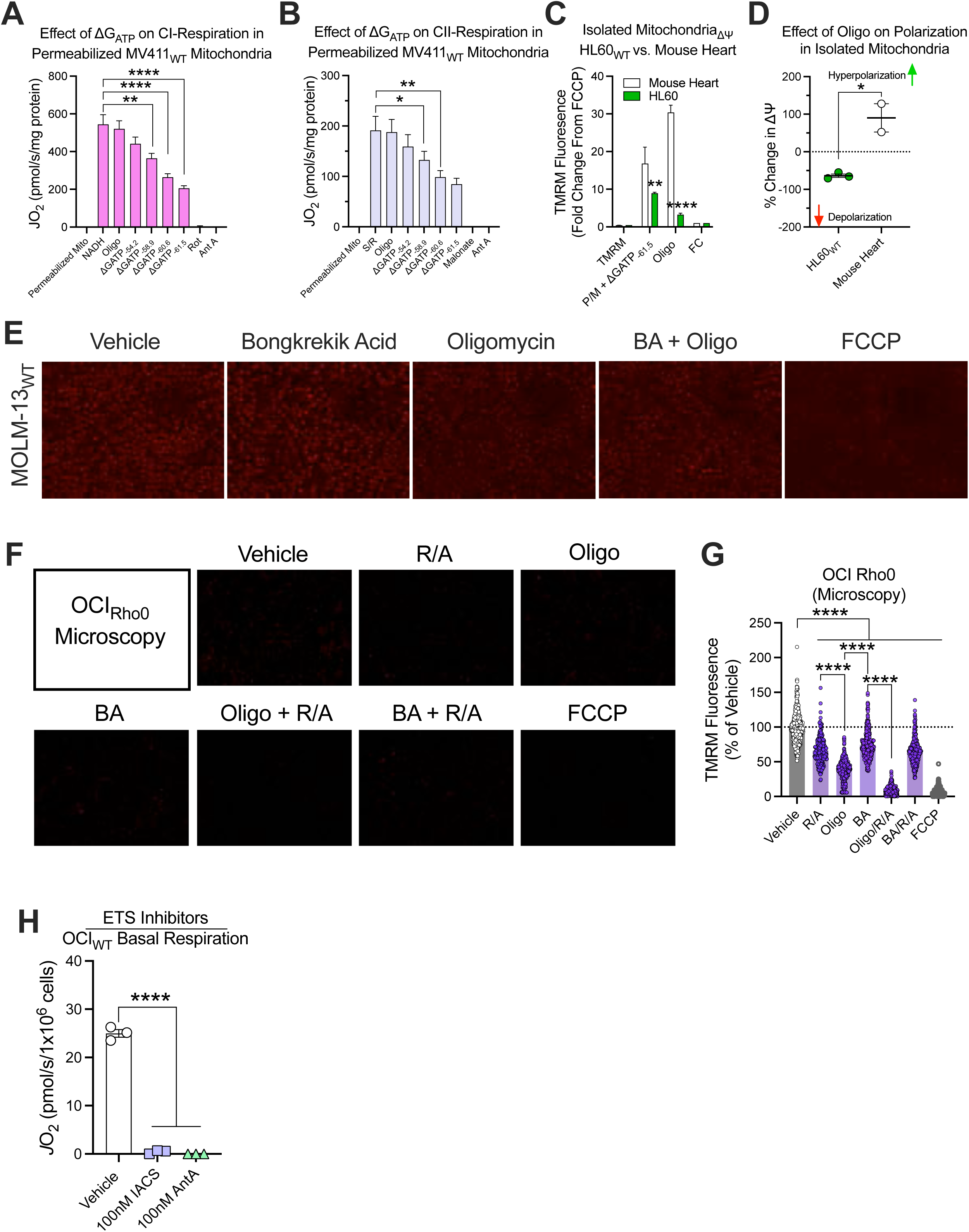
Extended validation of the potential mechanism of oligomycin-induced apoptosis. (Related to Figure 3) All experiments were performed using whole intact cells, digitonin-permeabilized cells, or isolated mitochondria. (A) Effect of ΔG_ATP_ on NADH (Complex I) supported respiration in alamethicin permeabilized isolated MV411_WT_ mitochondria (*n* = 4 replicates). (B) Effect of ΔG_ATP_ on succinate (Complex II) supported respiration in alamethicin permeabilized isolated MV411_WT_ mitochondria (*n* = 4 replicates). (C) Flow cytometric analysis of the ΔΨ_m_ of isolated mitochondria derived from mouse heart or HL60 cells (*n* = 2-3 replicates). (D) Comparison of oligomycin induced polarization of isolated mitochondria derived from mouse heart or HL60 cells (*n* = 2-3 replicates). (E) Fluorescent microscopy images of ΔΨ_m_ in intact MOLM-13_WT_ cells in the presence or absence of oligomycin, bongkrekic acid, or that in combination. (F) Fluorescent microscopy analysis of intact cell ΔΨ_m_ in OCI_Rho0_ cells in the presence of rotenone and Ant A, bongkrekic acid, and oligomycin as single agents, or rotenone and Ant A in combination with oligomycin or bongkrekic acid. (*n* = 251-435 cells). (G) Effect of IACS or Ant A on OCI_WT_ basal respiration (*n* = 3 replicates). Data are presented as mean ±SEM and analyzed by two-way ANOVA (C,G), one-way ANOVA (A,B,H), or unpaired t-test (D). *p<0.05, **p<0.01, ***p<0.001, ****p<0.0001.

**Supplementary Figure 4.**
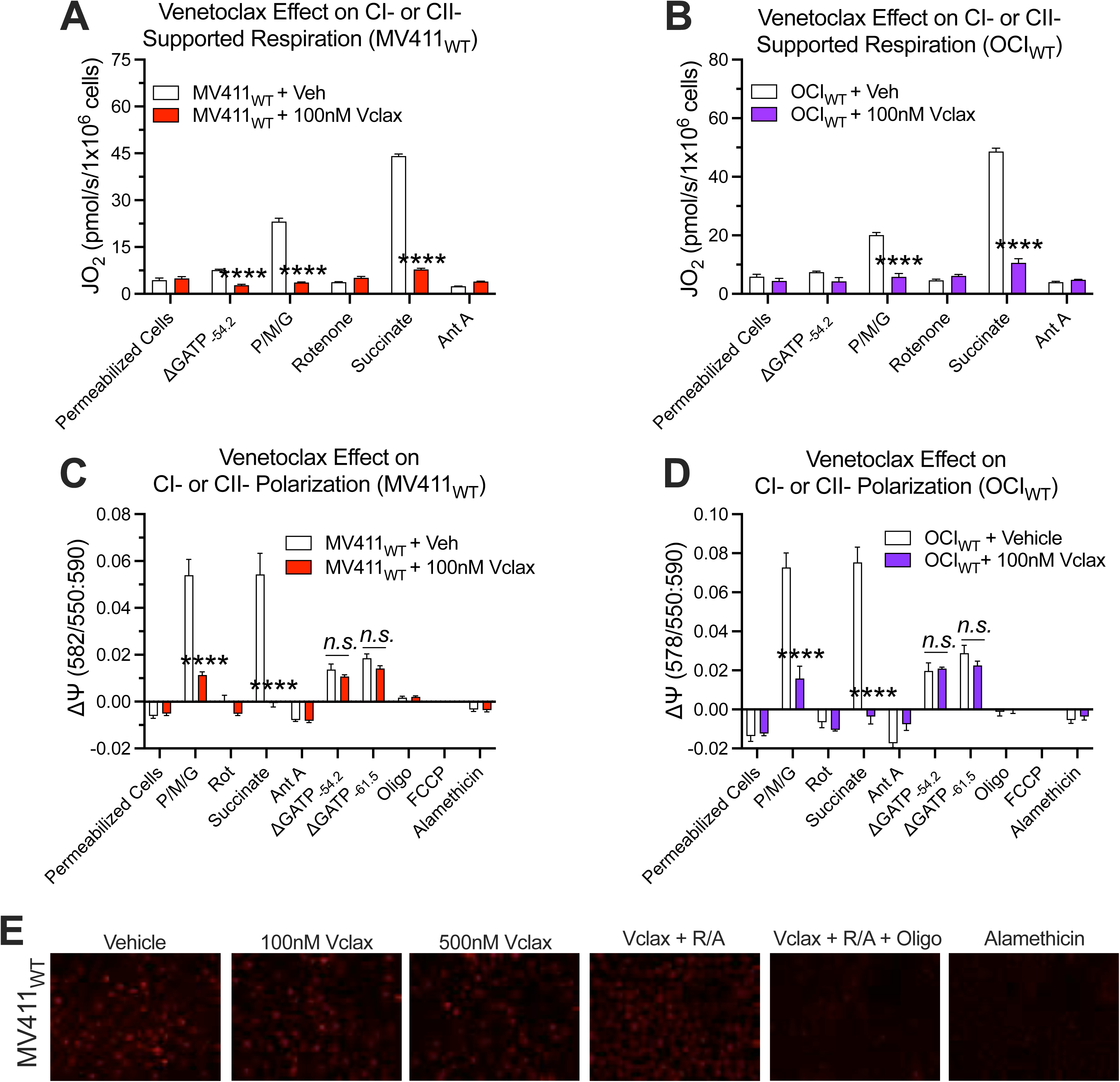
Complete and representative data for the experiments involving venetoclax exposure. (Related to Figure 4) All experiments were performed using whole intact cells or digitonin-permeabilized cells. (A) Effect of 1 hour exposure to 100nM venetoclax on respiration supported by Complex I substrates (P/M/G) or Complex II substrates (S/R) in MV411_WT_ cells (*n* = 4 replicates). (B) Effect of 1 hour exposure to 100nM venetoclax on respiration supported by Complex I substrates (P/M/G) or Complex II substrates (S/R) in OCI_WT_ cells (*n* = 4 replicates). (C) Effect of 1 hour exposure to 100nM venetoclax on polarization stimulated by Complex I substrates (P/M/G) or Complex II substrates (S/R) in MV411_WT_ cells (*n* = 6 replicates). (D) Effect of 1 hour exposure to 100nM venetoclax on polarization stimulated by Complex I substrates (P/M/G) or Complex II substrates (S/R) in OCI_WT_ cells (*n* = 4 replicates). (E) Fluorescent microscopy images of ΔΨ_m_ in MV411_WT_ cells exposed to venetoclax, venetoclax in combination with rotenone and antimycin A, or venetoclax in combination with rotenone, antimycin A, and oligomycin. Data are presented as mean ±SEM and analyzed by two-way ANOVA (A,B,C,D). *p<0.05, **p<0.01, ***p<0.001, ****p<0.0001.

**Supplementary Figure 5.**
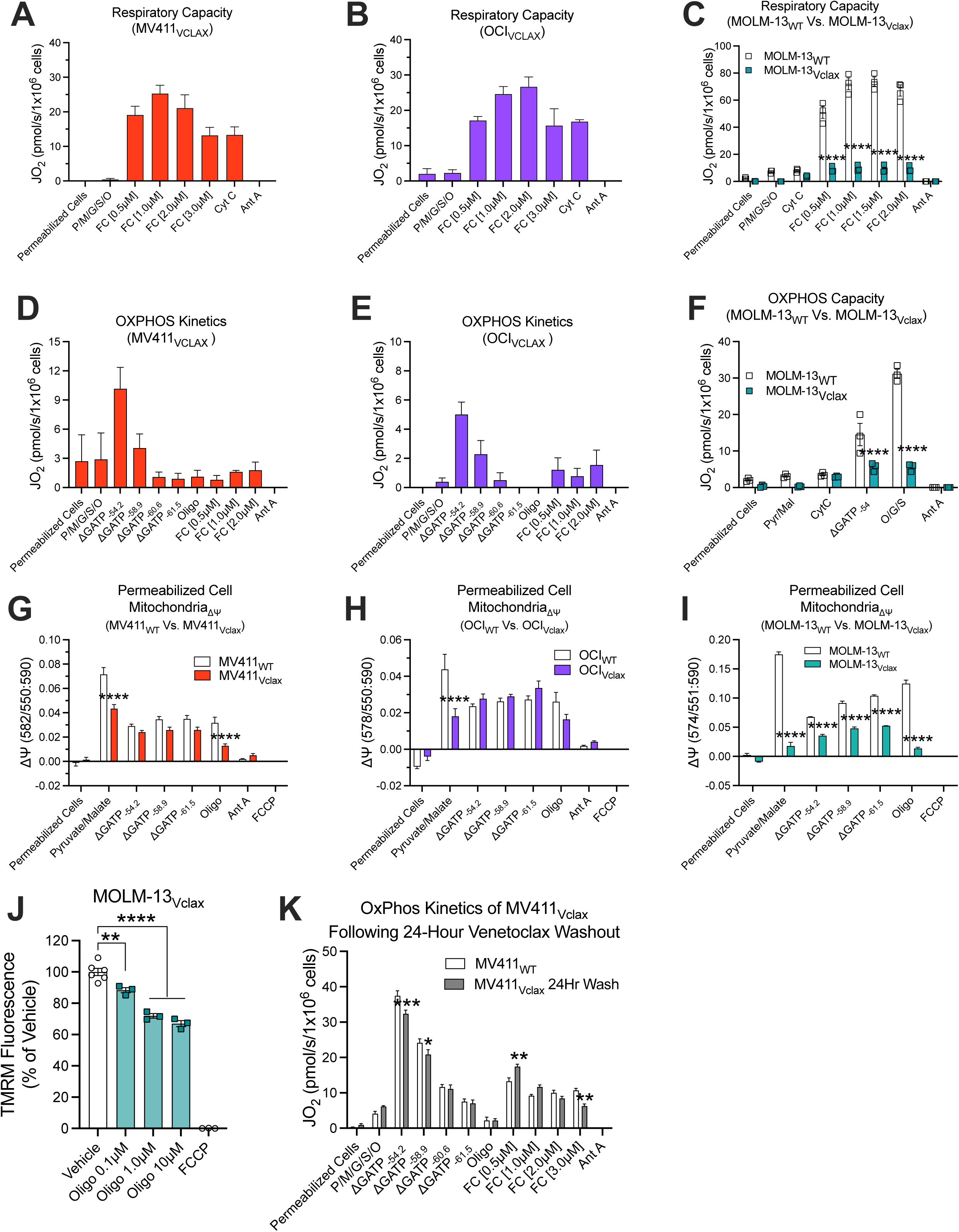
Complete and representative data for the experiments involving venetoclax resistance. (Related to Figure 5) All experiments were performed using digitonin-permeabilized cells. (A) Assessment of respiratory capacity in permeabilized MV411_Vclax_ cells (*n* = 3 replicates). (B) Assessment of respiratory capacity in permeabilized OCI_Vclax_ cells (*n* = 3 replicates). (C) Comparison of respiratory capacity in permeabilized MOLM-13_WT_ cells or MOLM-13_Vclax_ cells (*n* = 3 replicates). (D) Assessment of OxPhos respiratory kinetics in permeabilized MV411_Vclax_ cells (*n* = 3 replicates). (E) Assessment of OxPhos respiratory kinetics in permeabilized OCI_Vclax_ cells (*n* = 3 replicates). (F) Comparison of OxPhos capacity in permeabilized MOLM-13_WT_ cells or MOLM-13_Vclax_ cells (*n* = 3 replicates). (G) Comparison of the ΔΨ_m_ in permeabilized MV411_WT_ cells and MV411_Vclax_ cells (*n* = 5-6 replicates). (H) Comparison of the ΔΨ_m_ in permeabilized OCI_WT_ cells and OCI_Vclax_ cells (*n* = 5 replicates). (I) Comparison of the ΔΨ_m_ in permeabilized MOLM-13_WT_ cells and MOLM-13_Vclax_ cells (*n* = 3 replicates). (J) Flow cytometric analysis of ΔΨ_m_ in intact MOLM-13_Vclax_ cells in the presence of oligomycin (*n* = 3-6 replicates). (K) Comparison of OxPhos respiratory kinetics in permeabilized MV411_WT_ cells or MV411_Vclax_ cells after 24 hours of culture in venetoclax-replete media (*n* = 4 replicates). Data are presented as mean ±SEM and analyzed by two-way ANOVA (C,F,G,H,I,K). *p<0.05, **p<0.01, ***p<0.001, ****p<0.0001.

**Supplementary Figure 6.**
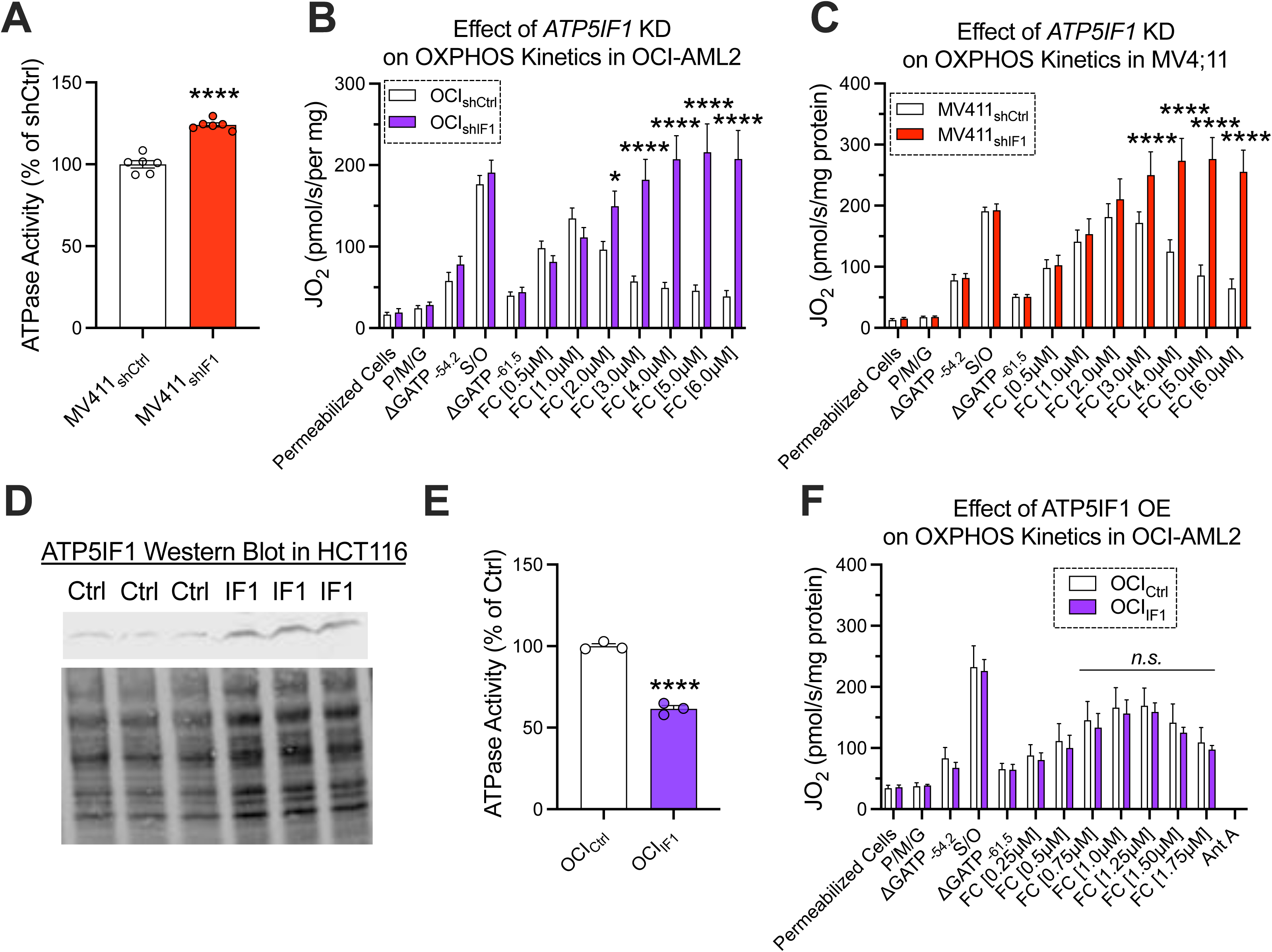
Profiling of OxPhos respiratory kinetics in *ATP5IF1* knockdown and overexpression AML cell models. (Related to Figures 6 and 7) All experiments were performed using digitonin-permeabilized cells. (A) Comparison of ATPase activity in isolated mitochondria lysates derived from MV411_shCtrl_ and MV411_shIF1_ cells (*n* = 6 replicates). (B) Comparison of OxPhos respiratory kinetics in permeabilized OCI_shCtrl_ cells or OCI_shIF1_ cells (*n* = 8-14 replicates). (C) Comparison of OxPhos respiratory kinetics in permeabilized MV411_shCtrl_ cells or MV411_shIF1_ cells (*n* = 9-10 replicates). (D) Western blot image of *ATP5IF1* expression in mitochondria isolated from HCT116_Ctrl_ and HCT_IF1_ cells. (E) Comparison of ATPase activity in isolated mitochondria lysates derived from OCI_Ctrl_ and OCI_IF1_ cells (*n* = 6 replicates) (F) Comparison of OxPhos respiratory kinetics in permeabilized OCI_Ctrl_ cells or OCI_IF1_ cells (*n* = 4 replicates). Data are presented as mean ±SEM and analyzed by two-way ANOVA (B,C,F) or unpaired t-test (A,E). *p<0.05, **p<0.01, ***p<0.001, ****p<0.0001.

**Supplemental Table 1.** Individual data points for all figures.

**Supplemental Table 2.** Proteomics Data. (**A**) Exported results from Proteome Discoverer 2.2 using the whole human proteome database. (**B**) Analyzed results, inclusive of Mitochondrial Enrichment Factor (MEF) calculation. (**C**) Exported results from Proteome Discoverer 2.2 using the human MitoCarta 3.0 database. (**D**) Analyzed results for MitoCarta 3.0 positive proteins.

